# Infection-Induced Extracellular Vesicles Evoke Neuronal Transcriptional and Epigenetic Changes

**DOI:** 10.1101/2020.11.26.399345

**Authors:** Ellen Tedford, Norhidayah Binti Badya, Conor Laing, Nozomi Asaoka, Shuji Kaneko, Beatrice Maria Filippi, Glenn Alan McConkey

## Abstract

Infection with the protozoan *Toxoplasma gondii* induces changes in neurotransmission, neuroinflammation, and behavior, yet it remains elusive how these changes come about. In this study we investigated how norepinephrine levels are altered by infection. TINEV (Toxoplasma-induced neuronal extracellular vesicles) isolated from infected noradrenergic cells down-regulated dopamine β-hydroxylase (*DBH*) gene expression in human and rodent cells. Here we report that intracerebral injection of TINEVs into the brain is sufficient to induce *DBH* down-regulation and distrupt catecholaminergic signalling. Further, TINEV treatment induced hypermethylation upstream of the DBH gene. An antisense lncRNA to *DBH* was found in purified TINEV preparations. Paracrine signalling to induce transcriptional gene silencing and DNA methylation may be a common mode to regulate neurologic function.

## INTRODUCTION

The mechanisms by which intracellular pathogens manipulate their host cells to change signalling pathways and the subsequent impact on host neuronal pathways requires delineation. *Toxoplasma gondii*, an intracellular protozoan in the phylum containing the malaria parasite *Plasmodium*, is a ubiquitous parasite that infects a wide range of warm-blooded animals with approximately one third of the global population seropositive (1). Seroprevalence in individuals ranges from 10-90% in different parts of the world with >40m people in the USA seropositive (DPDx, Centers for Disease Control, USA). Acute *T. gondii* infection is followed by a long-standing chronic infection of the brain and muscle that may last years, during which parasites are encysted as bradyzoite stages. In the brain, parasites are encysted exclusively in neurons. Inflammatory responses such as interferon gamma suppress active tachyzoite reproduction with major histocompatibility complex (MHC) I-dependent CD8+ T-cell recognition of infected neurons. Suppression of parasite growth in this way maintains the chronic bradyzoite infection in the brain (2–4). Changes in behaviour as well as neurotransmission and neurologic function have been observed during chronic infection and correlated with changes in dopamine, norepinephrine, glutamate and GABA. (5–13). In rodents, loss of innate fear and increased activity favour parasite transmission to the definitive feline host (14–16). Increased risk-taking and deficits in learning and memory have also been associated with *T. gondii* infection in animal models and human subjects (17–19). *T. gondii* seroprevalence and serointensity has been correlated with neurological disorders including depression, ADHD, epileptic seizures, movement disorders, and prominently with schizophrenia (3, 20–23).

The mechanisms responsible for behavioural and cognitive changes need delineation at the neurophysiological and gene expression levels. Recently, decreased norepinephrine (NE) concentrations and dopamine β-hydroxylase (DBH), responsible for its synthesis, have been observed in the brains of infected rodents and correlated with behaviour changes (7, 11). *DBH* expression was down-regulated 32 ± 2.1-fold compared to uninfected rats (p=0.0023) (7, 11). In that study, lower arousal measured as decreased locomotor activity at early time points (p<0.0001) was observed in infected mice and correlated with *DBH* expression. Increased sociability in infected mice, measured as social approach, also correlated with *DBH* expression. Arousal and sociability are noradrenergic-associated behaviours (7, 11, 24, 25). As NE also suppresses inflammatory cytokines (26, 27), decreases in concentration would augment the neuroinflammatory response (28). The observable changes in NE are surprising given the low percentage of cells infected. In a large study investigating brain cysts in rats, only 0.002-0.14% of cells contained tissue cysts for eleven *T. gondii* strains tested (29). In this study, we addressed the biological questions of the mechanism responsible for NE down-regulation and how *T. gondii* infection decreases brain NE levels when so few neurons contain the parasite. Initial experiments indicated a diffusible factor released by infected cells. EVs were isolated that induced transcriptional gene silencing (TGS) and chromatin remodelling.

## RESULTS

### Toxoplasma Infection Induces Release of Extracellular Vesicles from Host Cells that Decrease NE

Extracellular vesicles (EVs) have emerged as an important facet of host-pathogen interactions and were investigated for their relationship to the NE suppression (30, 31). EVs were purified from infected catecholamine-producing dopaminergic/noradrenergic cell cultures (henceforth termed noradrenergic cells) by step-wise ultracentrifugation followed by purification by sucrose-gradient fractionation (32, 33) (Fig. 1A). The EVs appear to be of host cell origin based on size, morphology, sedimentation rate, and exosome markers (34). The infected culture EVs contained mammalian exosomal protein markers CD81 and EpCAM (Fig. S1 and proteomic analysis, data not shown) as previously found with PC12 cells, as well as CD63, ICAM, FLOT-1, and TSG101 being detectable. The protein profiles were similar for EVs from infected and uninfected cultures (Fig. S1) and the size consistent with exosomes (Figs. 1B, 1C). Based on these properties, in this paper we have termed these *Toxoplasma-induced* neuronal host-derived extracellular vesicles (TINEV). Similar yields of released EVs were isolated regardless of infectious status in our experiments (5.6±1.4 μg/ml and 3.9±0.49 μg/ml, respectively; p=0.438) Infection naive noradrenergic cells were treated with TINEVs and expression of *DBH* was measured. Noradrenergic cells in medium with commercial exosome-depleted serum served as control. *DBH* expression was significantly down-regulated (190±67-fold, p=0.006 in Fig. 1D) in cultures treated with TINEVs.

**Figure 1:**
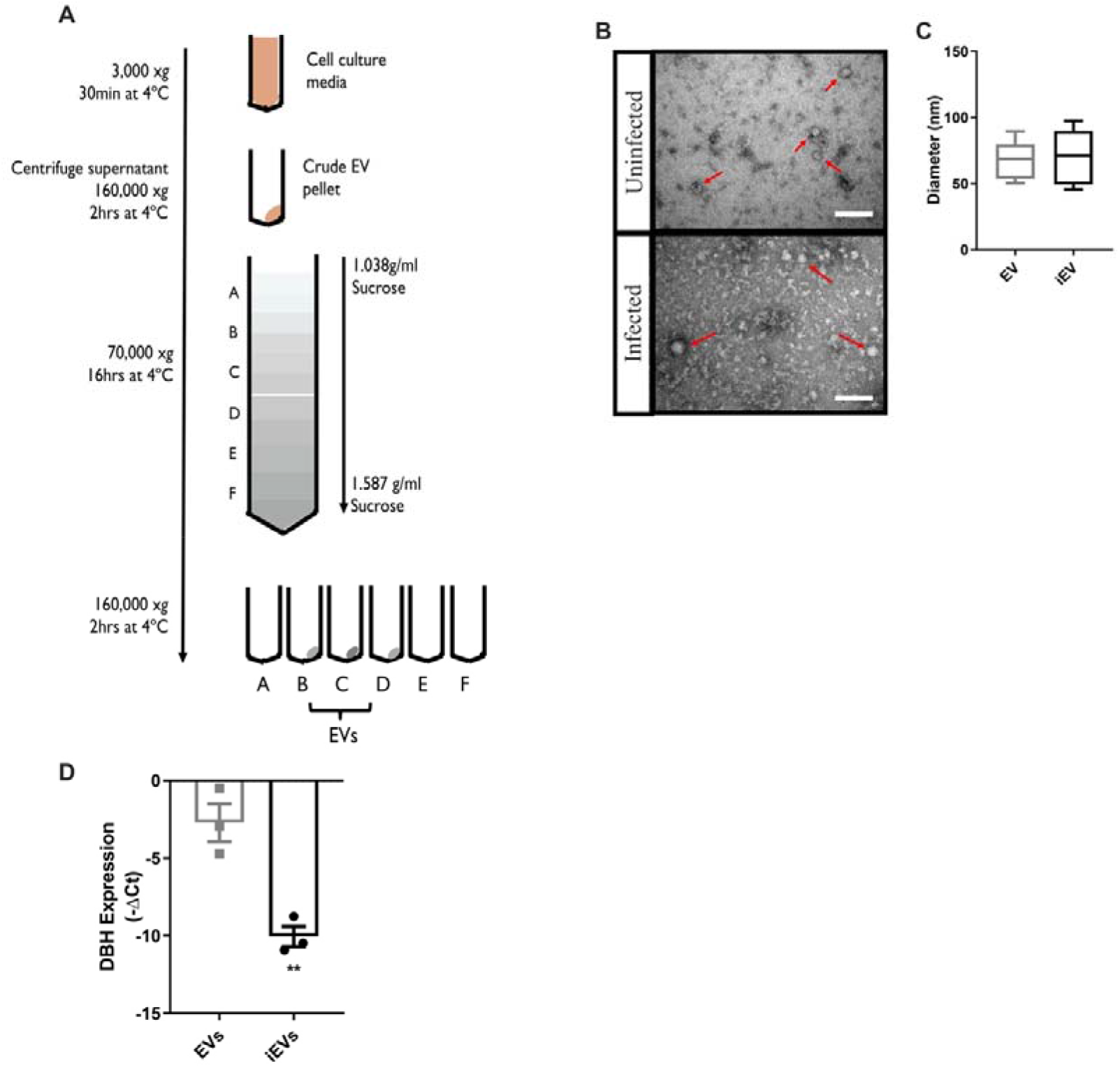
Toxoplasma-induced neuronal extracellular vesicles (TINEV) induce transcriptional regulation in neuronal cells. **A)** Scheme summarising sequential ultracentrifugation steps undertaken to produce purified extracellular vesicles. Sucrose gradient densities range from 0.104 g/ml to 1.43 g/ml. The density between 1.15 and 1.19 g/ml is collected as this represents extracellular vesicles (80). B) Transmission electron microscopy of extracellular vesicles isolated from uninfected (top) and infected (bottom) PC12 cells. Arrows indicate extracellular vesicles; scale bar represents 100nm. **C)** Plot of extracellular vesicle diameter from uninfected (EV) and *T. gondii* infected PC12 cells (iEV); p=0.81 **D)** *DBH* mRNA expression following treatment with purified TINEVs. PC12 cells were treated EVs from uninfected (grey) and *T. gondii*-infected (black) cells for 24 hours; n=3, ** p=0.006. Graphs show ±SEM with p values for Student’s t tests. E) A schematic representation of experimental design illustrating the site of injection for delivery of EVs directly into the brain’s noradrenergic center, the locus coeruleus (LC), of rats. Brain regions (prefrontal cortex, midbrain, and the pons/LC as illustrated by the dotted lines) were harvested two days following 3 days of twice daily treatment,. RNA was purified from tissues and RT-qPCR performed to quantitate *DBH* expression. Stereotaxic coordinates from Paxinos and Watson (81). **F)** *DBH* mRNA levels in brain tissues of rats receiving TINEVs (iEVs) and EVs from uninfected cells, relative to neuronal marker MAP2. The pons containing locus coeruleus (pons/LC) had significantly down-regulated *DBH*. ±SEM shown, n=5, Mann-Whitney unpaired t test, ** p=0.0079.

Next the *in vivo* effect of the TINEVs was examined. Intracranial injection of TINEVs into the *locus coeruleus* (LC), the central region of noradrenergic neurons in the brain, of adult rats was performed and *DBH* expression was measured by RT-qPCR (Fig. 1E, Fig. S2). TINEV injection induced a decrease in *DBH* mRNA (62±43-fold, p=0.0079) in the pons/LC region (Fig. 1F) compared to treatment with EVs from uninfected cultures. No difference was found in *DBH* mRNA levels in the mid-brain and prefrontal cortex for rats injected TINEVs, although low numbers of noradrenergic neurons are found in these regions. Health parameters (body weight, appearance and food intake) were normal in treated animals and expression of a housekeeping gene was unaltered in the brain with treatment (Fig. S3). Further, *DBH* expression in the adrenal glands was unaffected by the treatments (data not shown). The decrease in *DBH* mRNA observed with intracranial TINEV injection was similar to that observed in chronic infections (11).

Levels of NE and *DBH* mRNA have been found decreased with chronic *T. gondii* infection (11). A direct correlation between *DBH* expression and NE level (Fig. S4; correlation coefficient 0.81, p=0.014) was observed in rodent brains. Hence *DBH* expression can be used as a correlate of brain NE, as found in other studies (35). Based on the above findings, *T. gondii* infection induces cells to release TINEVs that are able to down-regulate *DBH* expression and cause a widespread decrease in brain NE.

### Strategy that Identified Transcriptional Gene Silencing and Epigenetic Change through Paracrine Signalling as the Mechanism of DBH Silencing

Initially, it was considered that several different factors could explain the disproportionate decrease in *DBH* expression (and hence NE) relative to the small percent of parasitised cells during chronic infection (e.g. neuroimmune responses). As decreased NE and *DBH* down-regulation (relative to other genes) has been observed *in vitro* with noradrenergic cell lines, mechanisms other than the host immune system are involved. A nuclear run-on assay was performed to assess whether the *DBH* down-regulation is at the transcriptional or post-transcriptional level. *De novo* transcription in nuclei isolated from infected cell cultures was measured by immunocapture of incorporated biotin-UTP. Lower amounts of nascent DBH mRNA (relative to standards) were found in host cell nuclei from infected than uninfected noradrenergic cell cultures (21±1.6-fold, p=0.00062) (Fig. 2A). Similarly, *de novo* transcription of *DBH* was downregulated in infected human noradrenergic cells (21±1.7-fold, p= 0.032) (Fig. 2B). Hence, infection induced transcriptional gene silencing (TGS) of *DBH* in human and rat noradrenergic cells.

**Figure 2:**
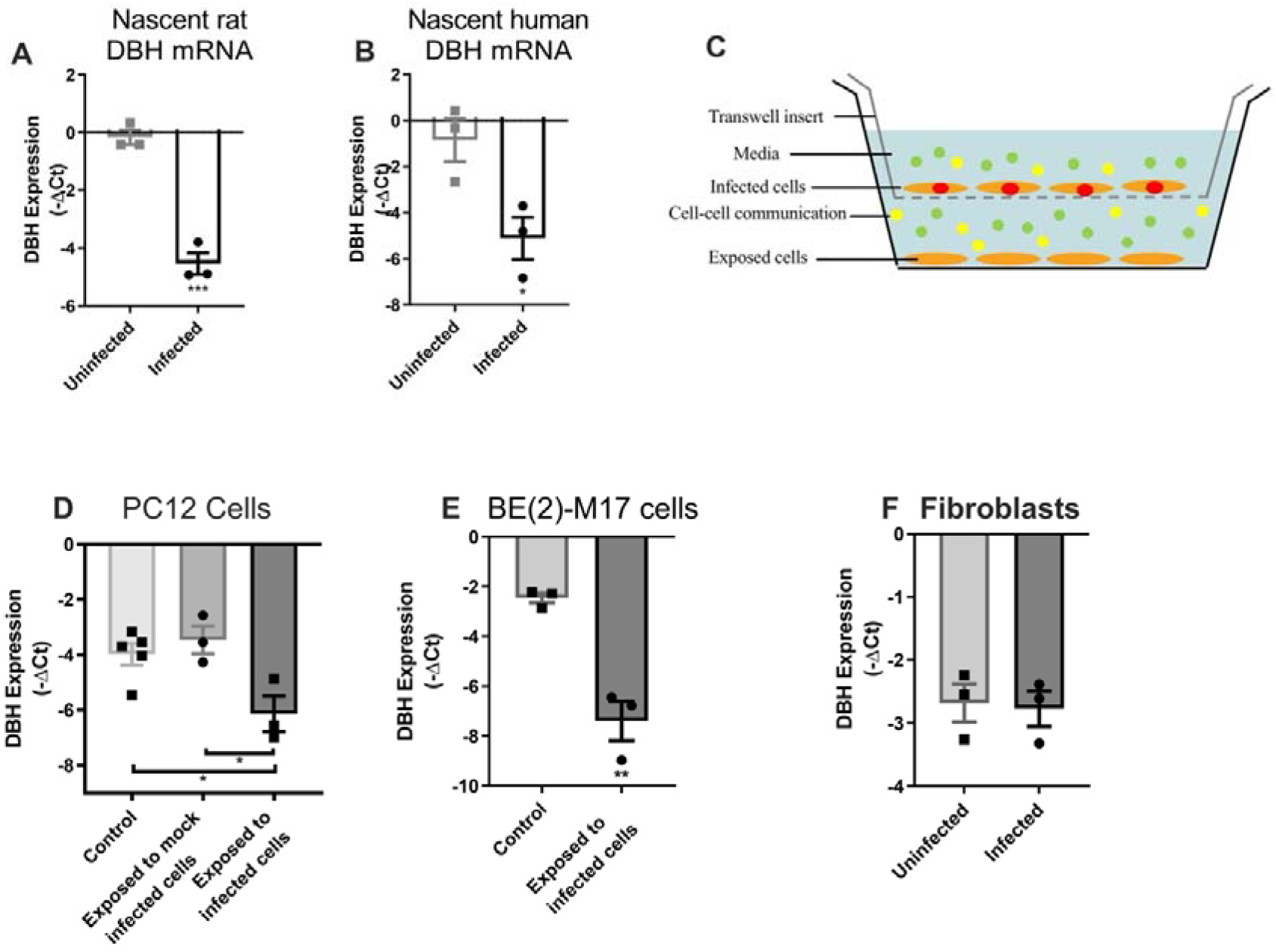
Noradrenergic cells infected with *T. gondii* secrete a signalling factor that transcriptionally regulates *DBH.* **A)** Nascent *DBH* pre-mRNA transcription (relative to GAPDH) in catecholaminergic PC12 cells infected with pH-shocked parasites. After 5 days of infection nuclear run-on assays were performed. MOI 1; n=3; ***p<0.001. B) Nascent *DBH* expression in uninfected and infected human neuronal cells as in (A). Multiplicity of infection (MOI) 1; n=3, *p<0.05. C) Scheme of transwell culture system with cells and *T. gondii* on the top layer and uninfected noradrenergic cells on the bottom layer separated by a 0.4μm membrane. Cells are orange, parasites red, signalling factors in vesicles (green) and soluble (yellow). **D)** *DBH* expression measured using RT-qPCR (relative to GAPDH) in PC12 cells from the bottom layer with either uninfected control, *T. gondii* infected, or mock-infected cells in the upper transwell layer. Mock-infected cells were incubated with heat-killed *T. gondii* tachyzoites n=3, * p<0.05. One-way ANOVA p= 0.017, Tukey’s post hoc test uninfected and mock-infected vs infected p=0.020 and p=0.033, respectively. E) Noradrenergic BE(2)-M17 cell expression of *DBH* mRNA cultured in transwells and treated as in (C); n=3, p=0.0038. F) Infected fibroblasts in the upper layer of the transwell system. *DBH* expression levels were measured in exposed noradrenergic cells. *DBH* gene expression was not significantly altered in cells exposed to infected fibroblasts (p=0.84); ±SEM shown, n=3, Student’s t test.

We examined the uninfected cells in cultures to determine whether those cells exposed to infected cells were also suppressed in NE as this could help explain the large change in expression observed. A transwell system was used that permits uninfected cells to be exposed to *T. gondii-ïnfected* cell products. This method was chosen because it differentiates between diffusible signals and parasites injecting components into cells without invasion, as was observed with a *Cre/loxP* assay in infected mouse brains (36). *DBH* expression was measured in uninfected rat noradrenergic cells in the bottom reservoir of the transwell system with the top reservoir containing an infected culture (Fig. 2C). *DBH* expression in the cells exposed to infected cultures was found to be down-regulated (7.9±2.8-fold, p=0.02) suggesting that a transmissible factor was released from infected cells that was subsequently identified as EVs. In contrast, noradrenergic cells that were exposed to cultures containing heat-killed *T. gondii* (‘mock-infected’) were unchanged in *DBH* expression (Fig. 2D). Exposure of a human neuronal cell line to infected cells in the transwell system induced a larger decrease in *DBH* expression (37±16-fold, p=0.0038) than the rat cell line (Fig. 2E). The observed *DBH* down-regulation is likely to be a minimal baseline as it remains possible that vesicles may stick to the transwell membrane and EV passage restricted. Parasite restriction to the upper reservoir of the 0.4 μm filter transwells was confirmed by inoculating standard *T. gondii* cultures with media removed from upper and lower reservoirs and monitoring propagation (data not shown). Transwells were set up with infected fibroblasts in the top reservoir and uninfected noradrenergic cells in the bottom reservoir to assess whether the down-regulation was cell-type specific (Fig. 2F). The noradrenergic cells exposed to *T. gondii-infected* fibroblast cultures were unchanged in *DBH* gene expression.

A further indication that EVs were the permeable effector responsible for the *DBH* down-regulation was finding that the insoluble components, separated from soluble factors by ultracentrifugation, contained the *DBH* down-regulating activity in preliminary tests (data not shown). This provided the rationale for EV isolation and testing.

### Epigenetic Changes with *DBH* Down-Regulation during Infection

As our findings indicated that TGS was responsible for *DBH* expression changes, the epigenetic state of the *DBH* gene was investigated (37). Methylation Sensitive Restriction Enzyme qPCR (MSRE-qPCR) was used to monitor DNA methylation levels in the *DBH* gene’s upstream region where the majority of CpGs are clustered (Fig. 3A). Methylation in the *DBH* upstream region rose from 16±4.6% to 66±3.8% in infected cultures of noradrenergic cells during the course of the infection (Fig. 3B; p=0.00072). As the noradrenergic cells are sensitive to pH changes (ie. neurotransmitter synthesis and synaptic transmission affected) (38, 39), alkaline-shocked tachyzoites were used for *in vitro* infections, as in prior studies (40). This procedure elevated expression of bradyzoite markers BAG1 and SAG4 (Fig. S5). *DBH* methylation was also increased in infected human noradrenergic cultures (2.8-fold; range 2.3-4.8-fold; p=0.0011) with a time-dependent increase in methylation in the region profiled (Fig. 3C).

**Figure 3:**
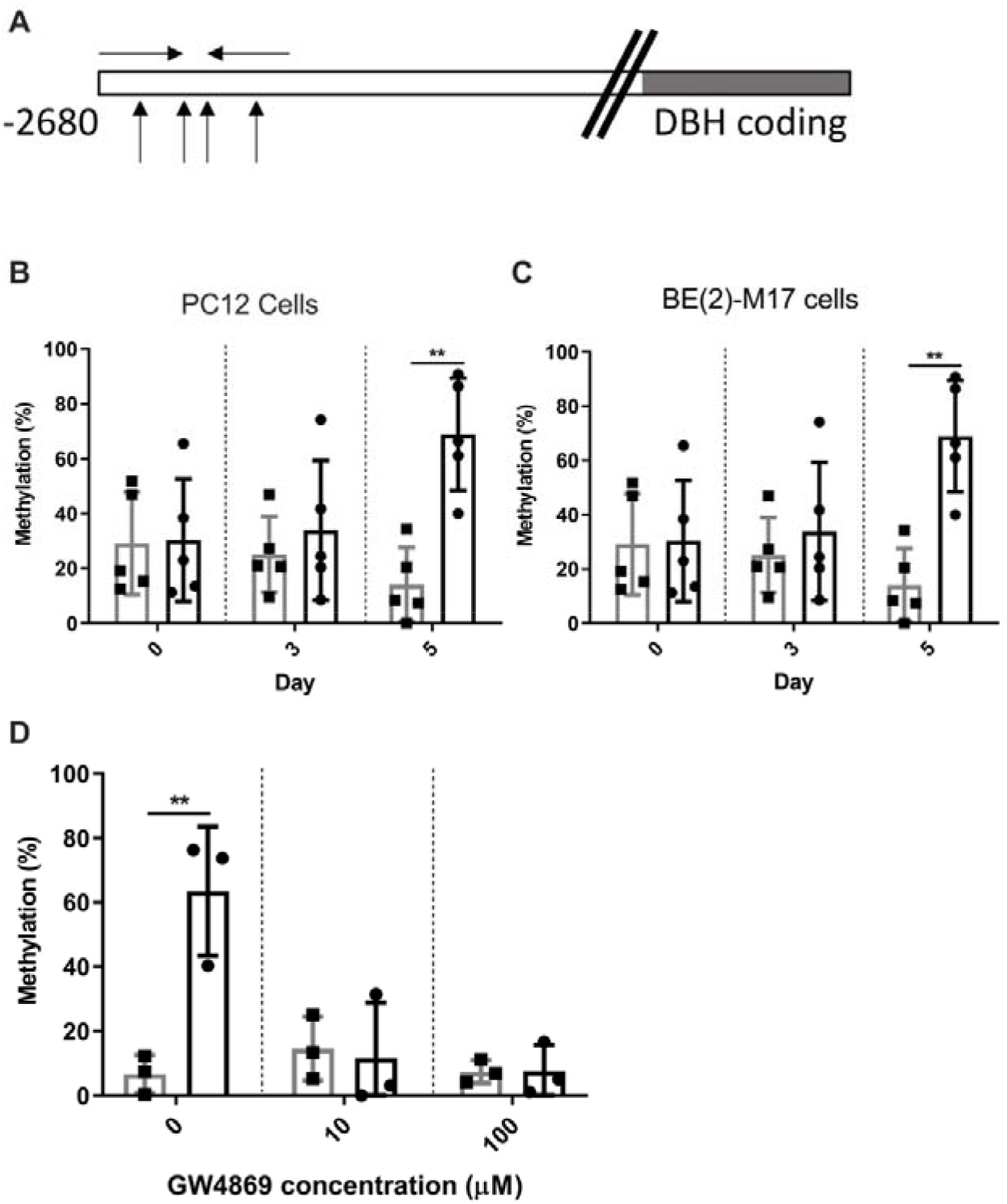
The promoter region of the DBH gene is hypermethylated during *T. gondii* infection. **A)** Diagram depicting the *DBH* promoter analysis by Methylation Sensitive Restriction Enzyme qPCR (MSRE-qPCR). The DBH coding region is shaded grey. The location of the methylation sensitive restriction enzyme sites are shown with up arrows and the primer sites with facing arrows (not to scale). **B)** *DBH* promoter methylation was measured by MSRE-qPCR. Percentage of methylation in the 5’ promoter region of the *DBH* gene in PC12 cells was measured by MSRE qPCR. DNA was collected at 0, 3 and 5 days post-infection from uninfected (grey) or infected (black) rat catecholaminergic cells. MOI=1; n=5, ***p= 0.0007. C) Methylation measured as in (A) during course of infection for human neuronal (BE(2)-M17) cells. MOI=1; n=5, **p= 0.0011. Unpaired Student’s t tests, ± SEM shown. D) Treatment with GW4869, a neutral sphingomyelinase (N-SMase) inhibitor, disrupted the DBH promoter methylation by permeable factors in the transwell system. DBH promoter methylation in the lower layer exposed cells was quantified by MSRE qPCR as described above. ± SEM shown, n=3, Student’s t-test, ** p=0.0093.

As an indicator of EV involvement in the epigenetic changes, cells were treated with an inhibitor of EV biogenesis in the transwell system that restricts parasites and parasite-infected cells from uninfected cell culture. GW4869 inhibits sphingomyelinase which is required for vesicle budding in endosomal formation. Addition of GW4869 to the transwell system abrogated the *DBH* hypermethylation observed in uninfected noradrenergic cells exposed to the infected culture (Fig. 3D).

In order to examine the epigenetic effects of *T. gondii* on *DBH* in neurons, *ex vivo* experiments were performed with infection of organotypic brain slices of the prefrontal cortex, nucleus accumbens and ventral tegmental areas and the TGS in the neuron population measured, as the percentage of infected neurons during chronic infection is 0.002-0.14% (29, 41). Methylation of the *DBH* upstream region rose from 29±2.7% in the brain tissue slices to 74±4.6% in the infected slices (p= 0.000051) (Fig. 4A). DNA methylation levels were then analysed in the brains of chronically-infected mice. Neurons were purified by FACS and *DBH* methylation measured. The *DBH* gene in infected animals was 53±7.7% methylated compared to 6.3±2.0% in uninfected mice (p= 0.000045, Fig. 4B). For comparison, levels of total genomic DNA methylation was measured (Fig. 4C). No change in global DNA methylation was observed, as has previously been found with *T. gondii* infection (42). Hypermethylation of CpG residues upstream of the *DBH* gene were also found in NGS genomic bisulfite sequencing of infected cultures (Fig. S6).

**Figure 4:**
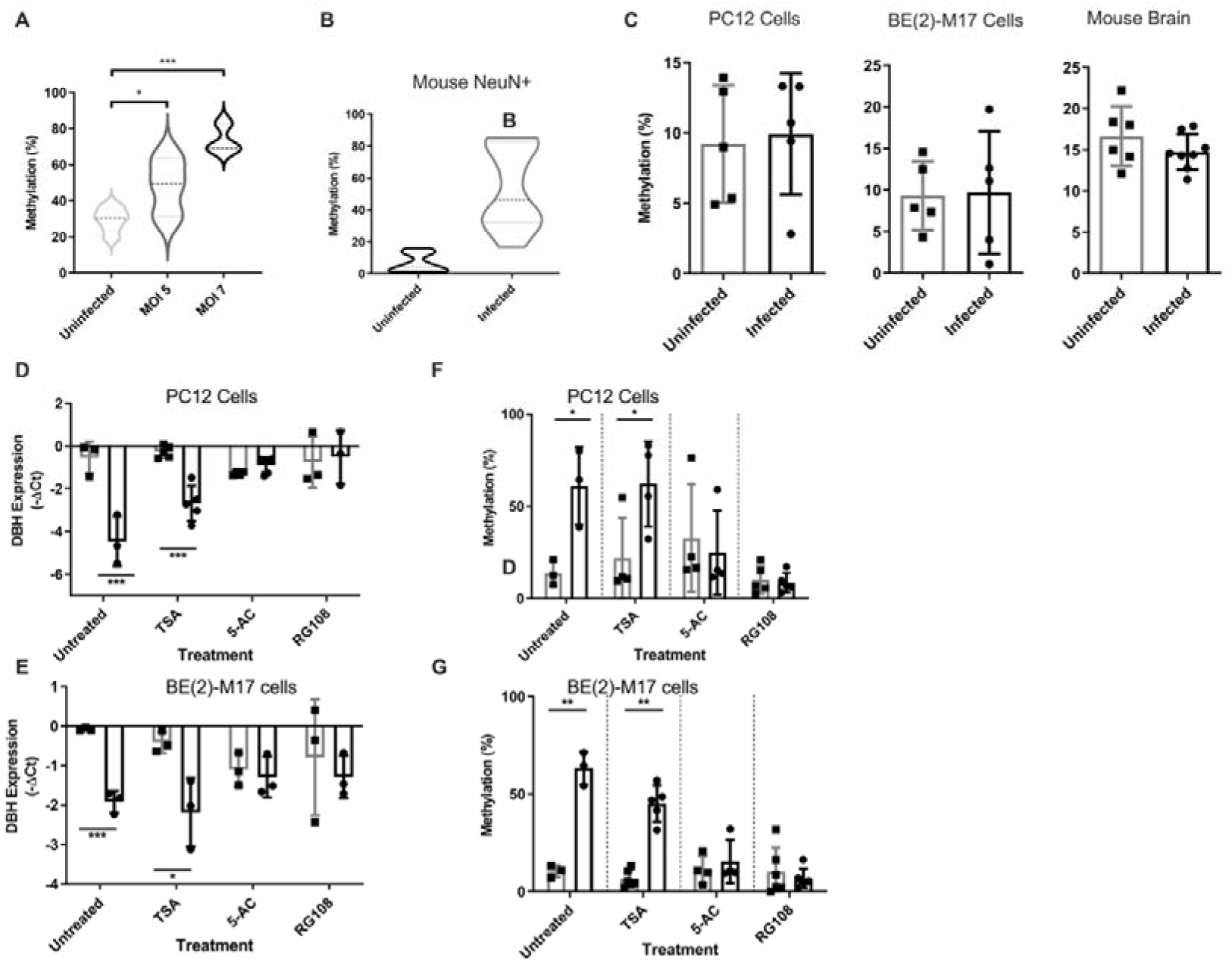
Epigenetic changes with gene expression of DBH with infection. **A)** *DBH* promoter methylation was measured by MSRE-qPCR in organotypic coronal midbrain section cultures. Slice cultures were either uninfected (light grey, n=6) or infected with wild-type induced-bradyzoite *T. gondii* at MOI 5 (dark grey, n=3) and MOI 7 (black, n=3). Infection was monitored for 5 days by light microscopy; n=3 biological repeats containing 3 slices from the same rat per well, * p=0.04, *** p= 0.000051 shown. Oneway ANOVA, p= 0.0005 Tukey’s post hoc test MOI 7 vs uninfected p=0.0004. **B)** Methylation at the *DBH* promoter region in neurons from uninfected (grey, n=9) and chronically-infected (black, n=11) mice. Neurons were enriched from harvested brains by FACS using NeuN antibody and *DBH* methylation quantified by MSRE-qPCR. p=0.0004. C) Total genomic DNA methylation measured by ELISA of both uninfected and infected rat catecholaminergic PC12 cells (n=3, p=0.52), human neuronal BE(2)-M17 cell (n=5, 0.59) and neuronal nuclei enriched by FACS from mouse brain tissue for uninfected (n=6) and chronically-infected mice (n=3, p=0.15). No significant change in methylation was identified by Student’s t tests, ± SEM shown. D) Uninfected (grey) and *T. gondii-infected* (black) noradrenergic PC12 cells treated with trichostatin A (TSA), 5-azacytidine (5-AC) and RG108; n=5, ** p<0.01. Expression of DBH mRNA measured by RT-qPCR relative to GAPDH. E) As in (A) with noradrenergic human BE(2)-M17 cells treated with TSA, 5-AC and RG108; n=5, * p<0.05. n=5. Student’s t test, ±SEM shown. **F)** 5’ DBH promoter hypermethylation was measured via MSRE qPCR (as above). Uninfected (grey) and *T. gondii-infected* (black) PC12 cells treated with trichostatin A (TSA), 5-azacytidine (5-AC) and RG108; n=5, ** p<0.01. **G)** Uninfected (grey) and infected (black) BE(2)-M17 cells treated as in (A) with TSA, 5-AC and RG108; n=5, * p<0.05. Student’s t test, ±SEM shown

The mechanism responsible for the parasite-induced DNA methylation and chromatin remodelling during *DBH* TGS was investigated. Experiments examined the role of DNA methyltransferase (DNMT) and histone deacetylation in this process. Infected noradrenergic cells were treated with the DNMT inhibitors RG108 and 5-azacytidine (5-AC). Both compounds disrupted the parasite-induced *DBH* down-regulation and DNA hypermethylation in rat noradrenergic cells, relative to marker (Figs. 4D, 4F). The DNMT inhibitors similarly blocked the TGS and epigenetic changes in human neuronal cells (Figs. 4E, 4G). This was not due to parasite sensitivity to the inhibitors since the inhibitors were non-toxic to parasites at concentrations tested (data not shown) and *T. gondii* lacks 5-methylcytosine (43). This implies that the *DBH* silencing involves DNMT. In contrast, the histone deacetylase inhibitor trichostatin A did not abrogate the *DBH* down-regulation or hypermethylation in infected cultures (Fig. 4D-G). Hence the mechanism may not involve histone deacetylation or activation was not captured within the experimental timeframe of 5 days and histone deacetylase is active at a different time in the epigenetic modification pathway.

To directly compare the epigenetic changes in infected versus uninfected (but infection-exposed) cells in the same culture, populations were enriched for cells containing GFP-expressing *T. gondii* (GFP+) and GFP-cells. *DBH* hypermethylation was found in both populations of cells (Fig. 5A, 5B; ANOVA test, p= 0.0089). Indeed, the GFP-cells exhibited DNA methylation levels at least equal to the GFP+ cells. Hence, we conclude that direct infection was not required for hypermethylation of the *DBH* gene in a cell.

**Figure 5:**
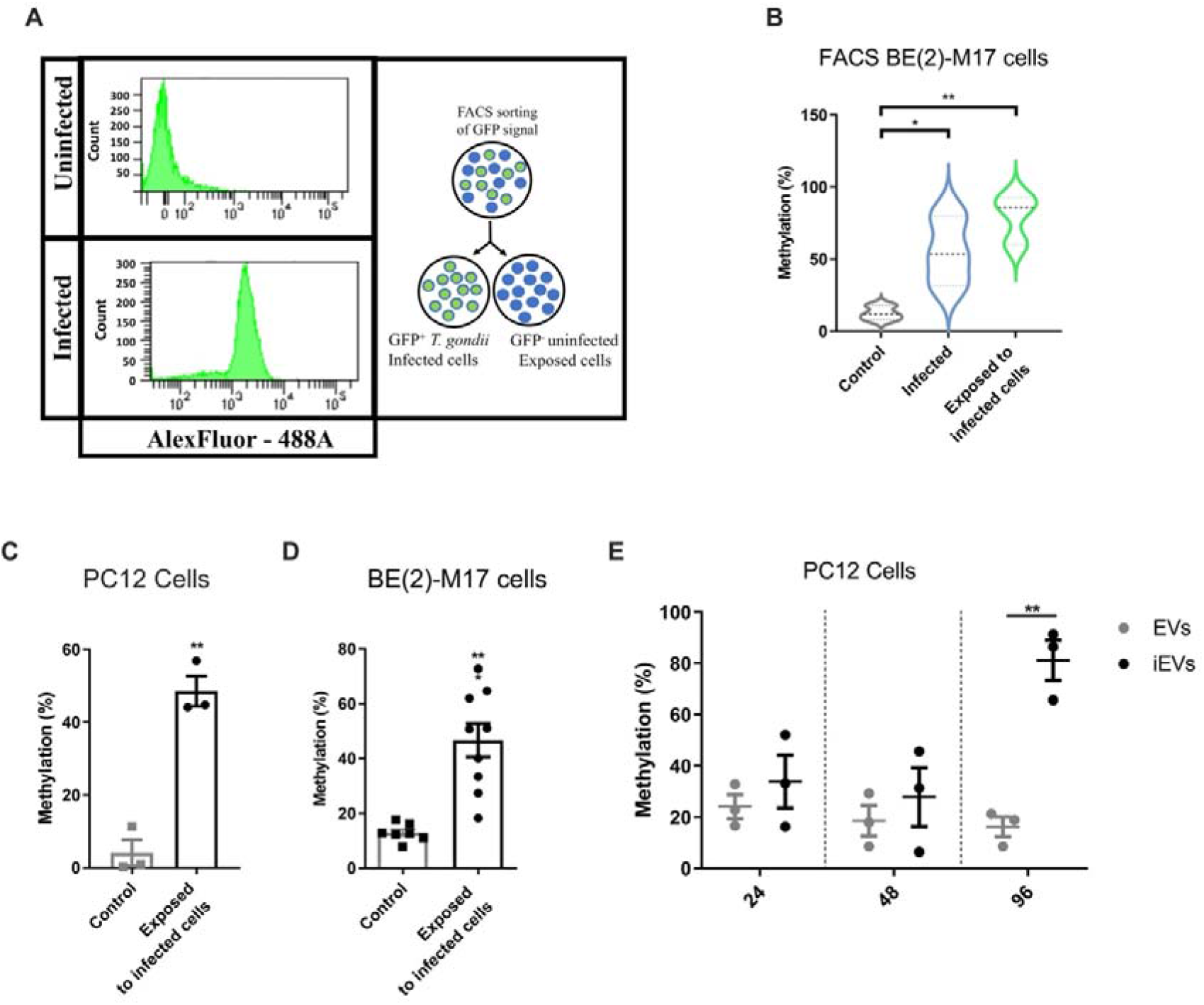
Hypermethylation of the *DBH* gene promoter region in uninfected, exposed cells. **A)** Representative plot showing cell count against GFP signal for FACs of GFP-negative and GFP positive cells infected with GFP-expressing parasites. Right panel shows a schematic representation of experimental design. B) Methylation at the 5’ prime *DBH* promoter region of uninfected cell control, infected GFP+, and exposed GFP-FACS-enriched BE(2)-M17 cells measured using MSRE qPCR; n=3, * p<0.05, ** p<0.01 shown. One-way ANOVA, p= 0.0089, Tukey’s post hoc test exposed to infected cells vs control p=0.0077. **C)** 5’ *DBH* region methylation in PC12 cells exposed to infected PC12 cells in a transwell system as Fig 2C. ±SEM shown, n=3, Student’s t test, ** p<0.01. **D)** BE(2)-M17 cell *DBH* promoter methylation measured as in (C) following 5 days transwell exposure to either uninfected (n=4) or infected BE(2)-M17 cells (n=9). ±SEM shown, Student’s t test, *** p<0.001. E) Time course of methylation changes induced by TINEV treatment. 5’ *DBH* methylation in cells treated with TINEVs (black) and EVs from uninfected cells (grey). Timepoints taken at 24, 48 and 96 hours of treatment; the minimum and maximum shown with line indicating mean; n=3, **p=0.0018. Graphs shown ±SEM with p values for Student’s t tests.

In order to identify the region of the *DBH* promoter involved in TGS in this model, the methylation status of the *DBH* upstream region in noradrenergic cells was measured by MSRE qPCR. Methylation of the *DBH* 5’ region was 34±4.8% greater (p=0.0013) in rat cells exposed to infected cells compared to exposure to uninfected cultures (Figure 5C).

Human noradrenergic cells exposed to infected cells also had increased methylation (49±1.9%, p=0.0003) of the DBH upstream region (Figure 5D). Hence, both transcriptional down-regulation and promoter hypermethylation are induced in human and rat noradrenergic cells exposed to *T. gondii* cell cultures.

Based on the above findings, the isolated TINEVs ability to induce epigenetic changes was next investigated. *DBH* gene promoter methylation, examined using the MSRE-qPCR assay, increased from 16±3.9% to 81±7.9% over the course of the experiment. In a time course of exposure to TINEVs, methylation was not significantly changed after 24 or 48 hours of exposure but hypermethylation was observed at 96 hours of incubation (Fig. 5E; ANOVA p=0.0008). Hence *DBH* transcriptional down-regulation and chromatin remodelling are induced by EVs from infected cells.

### Toxoplasma-induced Neuronal EVs Contain an Antisense DBH lncRNA

We investigated the molecular components involved in the TINEV-induced TGS and epigenetic modifications. Preliminarily, the sensitivity of the TINEVs to ultraviolet (UV) irradiation was tested to provide an indication of whether an RNA component could be involved (44). TINEVs treated with UV radiation no longer induced hypermethylation of the *DBH* promoter (Fig. S7). It is possible that UV radiation inactivated proteins.

With the specificity of the differential gene expression and the role of non-coding RNAs (ncRNA) in regulating gene expression, we explored the potential presence of a long non-coding RNA (lncRNA). LncRNA has roles in neuronal gene expression such as highly specific antisense transcripts (45–48). EVs have been found to contain miRNAs and lncRNAs, although their functional significance is still unclear. With the specificity of the *DBH* down-regulation in *T. gondii* infection and its magnitude with TINEV treatment, the identification of a long non-coding RNA (lncRNA) was investigated (45, 49). A panel of primers walking upstream stepwise from the *DBH* coding region were used to screen for an antisense RNA (37). These identified a lncRNA in infected cultures (Fig. S8). RNA purified from TINEV preparations were found to contain the *DBH* antisense lncRNA (Fig. 6A-C, Fig. S9). Although the lncRNA location as internal EV cargo is unconfirmed, it is unclear of the functional significance of the location (50). The lncRNA is complementary to the *DBH* upstream region containing cis-regulatory elements and crosses the transcription start site. As a positive control for samples, miR-21 miRNA served to confirm RNA quality purified from the TINEVs (51). Intriguingly, the timeframe observed for the *DBH* hypermethylation in this study (Fig. 5E) is similar to dynamic DNA methylation changes observed in cells treated with promoter targeted antisense RNA (52). The observations above represent the first study to show a specific and functionally-relevant lncRNA to be transmitted from one neuronal cell to another regulating neurotransmission (53).

**Figure 6:**
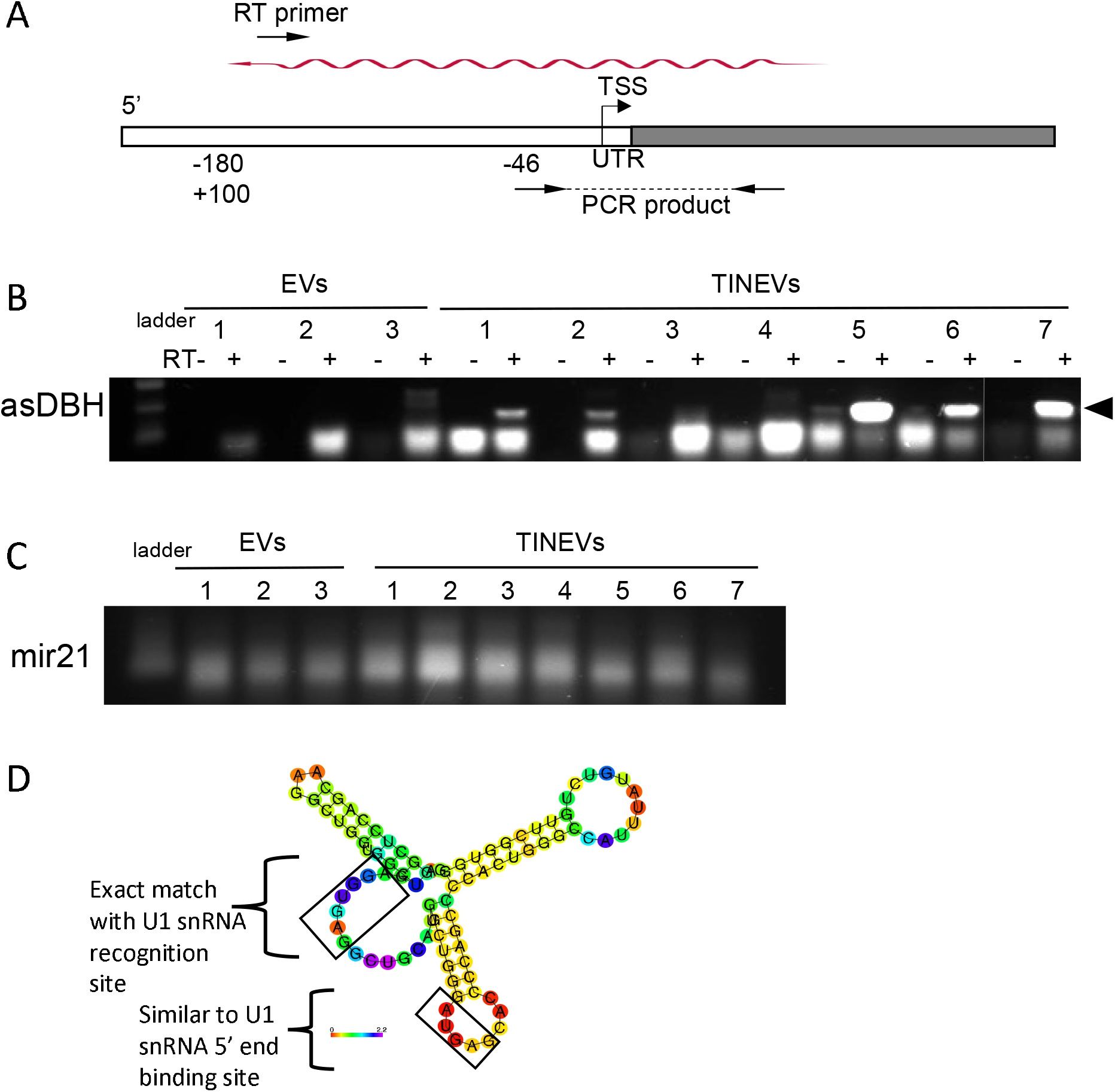
Toxoplasma-induced neuronal extracellular vesicles (TINEV) contain an antisense *DBH* lncRNA. **A)** Schematic representation of the *DBH* gene. The transcription start site (TSS) and untranslated region are highlighted with the binding sites of the directional RT and PCR primers used to detect the antisense lncRNA. The antisense lncRNA is depicted as the undulating arrow. **B)** Detection of asDBH by directional RT-PCR. TINEVs purified from repeated experiments (n=7) and EVs from uninfected cultures (n=3) resolved by gel electrophoresis with 1 kb Plus DNA Ladder (NEB). The product (arrowhead) is indicated (verified by sequencing). Alternating lanes are control reactions lacking reverse transcriptase. The full gel photo is shown in Fig. S9. **C)** An agarose gel of RT-PCR of miR-21 from all samples as a control for RNA quality. **D)** A drawing of the minimum free energy secondary structure prediction of asDBH using RNAfold with the base pair probabilities colored (82)and conserved binding sites for U1 snRNA identified as described in Yin et al (54).

The *DBH* antisense lncRNA was found to contain the conserved U1 snRNP recognition site that is a consensus sequence identified in chromatin-bound lncRNAs (54). Further, predicted secondary structure analysis places the U1 snRNP recognition site in a loop region in minimum free energy secondary structures of *DBH* antisense lncRNA (Fig. 6D). In the published study, the recognition site, importantly to be in a loop region, allowed binding of U1 snRNP to the lncRNA to promote chromatin interaction. Future experiments will further investigate this novel induced lncRNA.

## DISCUSSION

Intracellular infection can manipulate gene expression not only within the host cell but signalling to other cells. In this study it was found that *T. gondii* infection resulted in the release of EVs from noradrenergic cells that specifically induced gene silencing and DNA hypermethylation *in vitro* and in the brain. Many cells of the CNS release EVs including astrocytes, glia, oligodendrocytes, Schwann cells and neurons (55–57) and EV secretion is emerging as an important mechanism of cellular communication within the CNS (58). TINEVs purified from infected cultures and delivered into the brain induced *DBH* downregulation in neurons. EVs are continuously released in the brain limiting our ability to detect and isolate those induced by the 0.002-0.14% of cells that are infected during chronic *T. gondii* infection and restricted by the current lack of biomarkers for TINEVs. The findings presented here provide a mechanism that can resolve the enigma of an observable decrease in NE when only a small percentage of brain cells are infected. We propose a model of *T. gondii* inducing TINEV release with host lncRNAs that modulate the host environment (59).

Studies of extracellular vesicles secreted from pathogen-infected cells, including the apicomplexan parasites, have reported altering the host immune response (60–64). For example, cells infected with Epstein-Barr virus release EVs containing viral miRNAs that can downregulate cytokine expression in uninfected monocyte-derived dendritic cells. Human miRNAs complexed with Argonaute 2 are released in EVs from Plasmodium-infected red blood cells and induce pro-inflammatory cytokines in macrophages and endothelial cells. The findings reported here, in contrast, demonstrate a role for TINEVs in altering neurophysiology. TINEVs spreading to uninfected neurons carried host mammalian lncRNA, regulated gene expression, and modified chromatin. Intriguingly, NE modulates CNS cytokine release and hence the resultant NE decrease from *DBH* down-regulation will also affect the neuroimmune response. Host hypomethylation of the mammalian arginine vasopressin promoter in the amygdala of *T. gondii-infected* rodents has also been observed (42).

EV-miRNAs have been reported that alter DNA methylation in surrounding cells (63). Here, we identified a natural mammalian antisense lncRNA in TINEV preparations. The lncRNA is complimentary to the *DBH* 5’UTR and contains a consensus sequence for chromatin localization. It remains possible that *DBH* down-regulation is imposed by cell-autonomous immunity in noradrenergic cells by another component of TINEVs that has yet to be defined. Chromatin localisation of the lncRNA may occur via U1 snRNP binding at the transcriptionally active *DBH* gene and interacting with RNA polymerase II to disrupt transcription and facilitate DNA methylation. In this study, diffusible products of infected fibroblasts including EVs did not induce *DBH* down-regulation (Fig. 4F). In the future it will be interesting to assess the intercellular crosstalk from infected neurons to different brain cell types and to examine how *T. gondii* induces host cells to express and package selective lncRNAs.

Evidence is rapidly accumulating to support the role of EVs in the regulation of neurogenesis, neuronal connectivity and neuroimmune communication (44, 65). The findings in this study describe a novel form of neurotransmission regulation that may contribute to maintaining chronic infection. We identified a natural antisense lncRNA in TINEV preparations that specifically altered neurotransmitter signalling. In a recent study, reduced CNS noradrenergic signalling during chronic *T. gondii* infection correlated with noradrenergic-linked behaviour changes (11, 66). Chronic infection with bradyzoite-infected neurons permits continuous delivery of TINEVs into the brain. In this study, TINEVs delivered by intracranial injection twice daily for 3-days down-regulated *DBH.* As behaviour changes are measured in rodents during the 4th to 6th week of infection, a sustained supply of TINEVs (or the antisense lncRNA) is necessary for comparative behaviour tests. Also, reduced DBH has been found to elevate dopamine release from LC neurons (such as those modulating learning and memory) (7, 67) which could explain increased dopamine signalling that has been observed *T. gondii* infection (68). The neuroimmune system may also be affected by EVs. Changes in NE levels can alter cytokine responses via adrenergic receptors on astrocytes and microglia. Indeed, treatment with an adrenergic receptor agonist reversed the increased locomotion of infected mice (3, 69). Finding that EVs can modify DNA methylation of neurons may represent a mechanism, beyond parasitic machinery, in innate neurophysiological function and enhance our understanding of the role of dynamic DNA methylation in memory and learning.

## METHODS

### Cell culture

#### Parasite and Cell Culturing

*T. gondii* Prugniaud strain, isolated from mouse neuronal cysts, was used unless otherwise stated. Parasites were maintained in monolayer cultures of human foreskin fibroblasts (HFFs, ATCC® SCRC-1041) in Dulbecco’s Modified Eagles Medium (DMEM, GIBCO, USA) containing 10% foetal bovine serum (FBS) and 20 μg/ml^-1^ penicillin/streptmycin (Sigma, USA). Incubation of free tachyzoites in pH 8.2 DMEM with 5% FBS, overnight at 37°C was used to induce tachyzoites conversion to bradyzoites as previously described (11). Expression of *T. gondii* differentiation markers over 5 days of infection confirmed bradyzoite conversion (Figure S9). All experiments were performed with induced parasites unless otherwise stated.

Rat adrenal phaeochromocytoma, PC12, (ATCC® CRL-1721) cells were maintained at a density of 2-8×10^5^ cells/ml in Roswell Park Memorial Institute (RPMI) 1640 Medium (GIBCO, USA) and supplemented with 10% horse serum (GIBCO, USA), 5% FBS (GIBCO, USA) and 20 μg/ml^-1^ penicillin streptomycin (Sigma, USA). Human neuronal BE(2)-M17 cells (ATCC® CRL-2267) were maintained in a 1:1 ratio of F12 Hams and OptiMEM (GIBCO, USA) media supplemented with 10% horse serum (GIBCO, USA), 5% FBS (GIBCO, USA) and 20 μg/ml^-1^ penicillin streptomycin (Sigma, USA). The production of dopamine (DA) and norepinephrine (NE) by these cell lines was monitored regularly by HPLC-electrochemical detection. For experiments in which Extracellular vesicles (EV) were harvested or analysed, exosome depleted-FBS (GIBCO, USA) replaced the serum in the media for 24 hours prior to harvest. All cells were maintained at 5% carbon dioxide and 37°C and monitored for growth by light microscopy.

#### Infection of PC12 and BE(2)-M17 cells

Cells were cultured in multi-well plates at a density of 5×10^4^ cells/ml. When stated, transwell plates (Nunc™ cell culture inserts, Thermo Scientific) were used, with cells plated at 2.5 x 10^4^ cells/ml. Following 24 h of growth on 6-well plates, cells were infected with induced Prugniaud parasites in upper-wells, maintaining a multiplicity of infection (MOI) of 1. Cells were harvested immediately following infection (day 0) and after 3 and 5 days of infection for downstream processing. Mock-infection was performed using heat-killed *T. gondii* parasites subjected to 80°C for 10 minutes. The cultures were monitored daily by light microscopy.

#### Treatment of cells with purified TINEVs

EVs were extracted and purified from cell medium of uninfected and infected cells in exosome-depleted media, as described below. PC12 cells plated on 6-well plates and grown to a density of 2×10^4^ cells/ml were treated with EVs at a 10:1 culture media concentration equivalent (ie. 1 ml of PC12 cell culture treated with EVs purified from 10 ml of culture). Cells were treated every 24 h. DNA and RNA were harvested 24 h after the last treatment.

#### Drug assays

Cells were cultured and infected as previously described. Drug was added one hour before the time of infection (day 0). Cultures were grown in media containing a range of inhibitor concentrations; GW4869 (0-100 μM), Trichostatin A (TSA, 0-100 nM), RG108 (0-100 nM), or 5-Azacytidine (5-AC, 0-50μM) (SIGMA, USA). The effect of drug treatment on the parasites was monitored by infecting human fibroblasts with KU80-GFP bradyzoites or RH-YFP tachyzoites (70, 71) growth was measured by fluorescence using a FLUOstar Omega plate reader (BMG, UK).

#### Fluorescence-assisted cell sorting (FACS) of infected human neuronal cells

Human neuronal cells were infected with the Pru Δhpt/GFP/FLUC *T. gondii* strain, (kind gift from Boothroyd) (72). Five day infected and uninfected cell cultures were resuspended in PBS and separated in a CytoFLEX-2 flow cytometer (Becton Dickenson, USA) to isolate negative and GFP+ fluorescent cells. Control uninfected cell cultures were also subjected to FACS. DNA was extracted and purified from FACS populations.

### Animal Work

#### Organotypic brain slice cultures

All animal care and experimental procedures were approved by the Kyoto University Animal Research Committee. These experiments were performed with the help and guidance of the Kaneko Laboratory at Kyoto University. Sectioning was performed as described (41). Sprague-Dawley rat pups at postnatal days 3-4 were anesthetized by hypothermia. Brain was removed and separated into two hemispheres. Coronal sections (350μm thickness) were prepared under sterile conditions. Slices were dissected to include the ventral tegmental area (VTA). These slice cultures were placed on 30 mm insert membranes (Millicell 0.4μm; Millipore). Slice culture medium contained RPMI supplemented with 10% horse serum (GIBCO, USA), 5% FBS and additional 6.5 mg/ml glucose and 2 mM l-glutamine. The brain slice cultures were cultured at 37°C in a 5% CO^2^ for 15 days after dissection prior to use in experiments.

#### Rodent Infections

Brain samples were supplied from animal experiments conducted at the University of Leeds research animal facility with procedures approved by the University Animal Care and Use Committee and following Home Office, HSE, regulations and guidelines for Animals (Scientific Procedures) Act 1986 published in 2014 and with considerations of the replacement, reduction, and refinement in the use of animals for research.

#### Mice

Mice were housed up to five to a cage and maintained on a standard light-dark cycle with access to chow diet and water *ad libitum.* Throughout the experiments animals were monitored for illness or weight loss (more than 25%) and sacrificed 5-6 weeks after infection. Mice used were BALB/cAnNCrl x C57BL/6NCrl) F1 generation male mice. The mouse infections, brain dissection and frozen cryostat sectioning was performed as described (11).

#### Rats

All rat experiments were undertaken as previously described (11) prior to tissue sampling. Male Lister Hood rats (Harlan Ltd, UK) were used for all experiments. Rats were housed in individual cages and maintained on a standard light-dark cycle with access to chow diet and water *ad libitum.* Animals were infected by intraperitoneal (IP) injection with tachyzoites. Throughout the experiments animals were monitored for illness or weight loss (more than 25%) and sacrificed 5-6 months post-infection. Brains were snap-frozen for cryosectioning and RNA extraction.

#### Extracellular vesicle intracerebral treatment

Intracerebral injection into adult rats followed previously described experimental procedures (73). TINEVs extracted and purified from rat catecholaminergic PC12 cells as described below were used to treat rats. Eight-week-old male Sprague-Dawley rats weighing between 260 and 280g (Charles River Laboratories) were used. Rats were housed in individual cages and maintained on a standard light-dark cycle with access to chow diet and water *ad libitum.* Rats were stereotactically (David Kopf Instruments) implanted with indwelling bilateral cannula targeting the locus coeruleus. Health status was monitored visually and by food consumption (appetite) and body weight After one week of recovery, implanted rats received a 2 μl infusion twice a day for 3 days with TINEVs (4 μg protein) isolated from infected PC12 cells. EVs from uninfected PC12 cells served as a control. On the fourth day rats were anaesthetized and received an injection 3 μl bromophenol blue through each side of the bilateral DVC cannula (to verify their placement) prior to harvesting brain tissue. Brain tissue was dissected into prefrontal cortex, midbrain, and hindbrain regions for processing for RNA.

#### Fluorescence-assisted cell sorting (FACS) of mouse neurons

Mouse brain samples were processed as described (74). Briefly, brains were suspended in lysis buffer (0.32M sucrose; 0.1mM ethylenediaminetetraacetic acid; 5mM CaCl_2_; 3mM Mg(Acetate)2; 10mM Tris-HCl pH 8; 1mM dithiothreitol; and 0.1% Triton X-100) and processed using a 15 ml glass dounce homogenizer (Wheaton, UK) in 5-10 strokes on ice. Samples were centrifuged at 100,000xg with a two-step sucrose gradient to purify nuclei. Nuclei were stained with primary antibody NeuN (Abcam, EPR12763 1:250) and secondary antibody Alexa-488 (Abeam, ab150077,1:1000). All nuclei were also stained with Hoechst 33342 (SIGMA, USA). Samples were sorted using the Becton Dickinson FACS Aria II system.

### Extracellular vesicle characterization

#### Extracellular vesicle purification

TINEVs were isolated from media by ultracentrifugation as described (75, 76). For all experiments, EVs were isolated in parallel from uninfected cells. Briefly, one day prior to harvest, complete media was replaced with media containing exosome-depleted FBS (Systems Bioscience, CA). Cell cultures of uninfected and infected cells (five days following infection) were harvested and centrifuged at 3000xg for 10 minutes at 4°C. The supernatant was isolated and centrifuged at 160,000xg for 2 hours in a Type 60 Ti fixed angle rotor at 4°C. The pellet was resuspended in one ml of 90% sucrose. Eleven layers of one ml of sucrose 70%-10% w/v were then added and centrifuged at 70,000xg for 16 hours at 4°C in a Type 60 Ti fixed angle rotor. Supernatant was collected in two ml fractions. PBS (8 ml) was added to the fraction 1.1-1.2 g/ml sucrose with a density corresponding to that of EVs and centrifuged at 160,000xg for 70 minutes at 4°C to recover purified EVs. Methods and data on purification and characterization were input to the EV-TRACK knowledgebase (EV-TRACK ID: EV220363) (https://evtrack.org/).

#### UV Treatment

EVs were isolated as described. Prior to UV treatment, *DBH* down-regulation by TINEVs was verified following incubation of noradrenergic cells for 24 hours. TINEVs were exposed or mock exposed to UV light at 302nm for 5 minutes at 4°C using a transilluminator (Syngene, Cambridge).

#### Dot blotting

Dot blotting was performed using the Exo-Check Exosome Antibody Array (Systems Bioscience, CA) as per manufacturer’s instructions. Briefly, 500 μg protein equivalent of purified EVs isolated from PC12 culture were lysed and ligated to horseradish peroxidase (HRP) overnight at 4°C. Ligated protein was then incubated with the antibody membrane for two hours at room temperature. After three washes SuperSignal West Femto Chemiluminescent Substrate kit (Thermo Scientific, UK) was used as per manufacturer’s instructions. The blot was visualised via 90 second exposure to X ray film. Film was developed using the Konica SRX-101A Tabletop Processor.

#### Transmission electron microscopy (TEM)

EVs were purified as described above. Electron microscopy was performed as described (77). Freshly isolated EVs were resuspended in cold Dulbecco Phosphate-Buffered Saline (DPBS) containing 2% paraformaldehyde. EVs were mounted on copper grids, fixed with 1% glutaraldehyde in cold DPBS for 5 minutes at room temperature, negative stain was performed with uranyl-oxalate solution at pH 7 for 5 min, and embedded with methyl cellulose-UA for 10 min at 4°C. Excess cellulose was removed, and samples were dried for permanent preservation. Electron microscopy was performed in the University of Leeds facility using a Titan Krios 2 electron microscope. Analysis and extracellular vesicle diameter were measured using ImageJ software.

### Molecular Techniques

#### DNA Extraction

Classic phenol-chloroform method extraction was performed as described in (78). Briefly, tissue samples were first homogenised using a Micro Tissue Homogenizer (Fisher Scientific, UK). Homogenised tissue or cell pellets were incubated at 56°C overnight with 20 mg/ml Proteinase K. An equal amount of phenol/chloroform/isoamyl alcohol (PCI) solution (25:24:1) was then added, mixed and centrifuged. The aqueous layer was isolated and an equal volume of chloroform:isoamyl (CI) alcohol (24:1) added. DNA was precipitated using absolute ethanol and DNA, then resuspended in deionised water.

RNA extraction was performed using Direct-zol (Zymo, USA) as per manufacturer’s instructions.

#### PCR

Polymerase chain reaction was performed using the GoTaq PCR master mix (Promega, USA) as per manufacturer’s instructions. Samples were amplified in a 2720 thermal cycler (Applied Biosystems, USA). The PCR parameters were 94°C for 30 seconds; 35 cycles of 94°C for 30 seconds (denaturing step), 55°C for 15 seconds (annealing step) and 72°C for 10 seconds (elongation step); a final elongation of 72°C for 30 seconds.

Samples were visualised on a 2% w/v agarose gel with a Trackit 1kb Plus DNA ladder run at 80V until the DNA ladder and the sample products of interest had separated adequately.

#### RT-qPCR

RT-qPCR was performed using SYBR green master mix (Life Technologies, USA) as per manufacturer’s instructions and run using a CFX Max Real Time PCR machine (Bio-rad, USA). Parameters were 95°C for 2min and followed by 45 cycles of 95°C for 15 seconds, (primer Tm-5) °C for 15 seconds, and 72°C for 10 seconds. Primer sequences are recorded in Table S1.1. A melt-curve was performed to check for expected products. All RT-qPCR was performed with four technical replicates which were averaged.

#### Nuclear run-on

Nuclear run-on was performed as described (79). Briefly, after five days of infection, cells were incubated with lysis buffer containing 10 mM Tris-HCl pH 7.4,10 mM NaCl, 3 mM MgCl2, 0.5% NP-40 at 4°C isolating nuclei. Nuclei are then suspended in transcription buffer containing 20 mM Tris-HCl pH 8.3, 5 mM MgCl2, 300 mM KCl,4 mM DTT, RNase OUT 40 Units/μl, 0.5 mM BrUTP, 1mM ATP, 1mM GTP, 1mM CTP, 1mM UTP. Samples were incubated for 45 min at 37°C prior to RNA extraction using MEGAclear Transcription Clean-Up Kit (Life Technologies) as per manufacturer’s instructions. Bead-based immunoprecipitation was performed with protein G Dynabeads and IgG anti-BrdU antibody IIB5 (Santa Cruz Biotechnology, SC-32323). RNA was extracted and RT-qPCR performed as previously described.

#### Methylation sensitive restriction enzyme (MSRE) quantitative PCR

DNA was extracted from cell cultures and isolated as previously described. DNA concentrations were confirmed using a Nanodrop spectrophotometer. Samples were divided into reference and test samples. Test samples were digested with the high-fidelity methylation sensitive restriction enzymes (MSRE) HpaII, MaeII and SmaI (TaiI) in the Tango buffer (Thermo Fisher Scientific, USA). These MSREs were chosen based on the restriction enzymes sites present in the DBH upstream region. Reference samples were mock digested. Digestion was performed by heating to 25°C for 30 minutes, 37° for 30 minutes and 65° for 30 minutes. RT-qPCR was then performed. Percentage methylation calculated based on the difference in Cq values between test and reference samples. Digestion was confirmed with methylated and unmethylated human DNA standards provided with the OneStep qMethyl kit (Zymo) as per manufacturer’s instructions. Additionally, primers and MSRE digestion was confirmed with MSRE-PCR. qPCR was performed using SYBR®Green Real-Time PCR Master Mix (Thermo Fisher).

#### Directional RT-PCR for DBH lncRNA Screening

Three single-stranded forward (ie. sense) directional primers were designed to detect antisense RNA and are labelled RT-Primer 1, RT-Primer 2, and RT-Primer 3 (Extended Data Fig. 6A). The RT-designed primers are located (1) in the first exon, (2) near the transcription start site, and (3) −378 bp upstream of the DBH coding sequence. Four pairs of PCR primers were then designed to amplify products from the RT primer synthesized-template to screen for the presence of antisense lncRNA (Extended Data Fig. 6A). A list of primers used is shown, Table S1.1.

Total RNA was extracted from *T. gondii* infected-PC12 cells that were harvested on day five post-infection using the Direct-zol RNA MiniPrep Plus kit (Zymo Research, USA) according to the manufacturer’s instructions. DNase I treatment was implemented as suggested by the manufacturer. In addition, to ensure that all trace amounts of DNA contamination were removed, the eluted RNA samples were treated again with DNase enzyme using TURBO DNA-free™ kit (Invitrogen, USA) according to the manufacturer’s instructions as in earlier studies (37). The dependence of PCR products on the RNA samples was verified by parallel experiments with RNase treatment (data not shown). RNA samples were reverse transcribed with first strand synthesis primed with RT-primer 1 to 3 with a Maxima H Minus First Strand cDNA synthesis kit (Thermo Scientific, USA) following the manufacturer’s instructions. For each of the samples, there were several controls including priming the cDNA synthesis with random hexamer primer, a negative control reaction to assess the gDNA contamination in the RNA sample containing all components for RT-PCR except the Maxima H Minus enzyme mix, and no template control to assess reagent contamination. The mixtures were incubated as follows: 25°C for 10 min, 50°C for 30 min, 65°C for 20 min and then the reaction was terminated by heating at 85°C for 5 min. The reaction products served as templates for PCR with GoTaq ® G2 Hot Start Master Mix (Promega, USA), 300 nM forward primer, 300 nM reverse primer, and 2 μl template DNA. Thermal cycling was 3 min at 95°C, followed by 30 cycles of 95°C for 30 seconds, 57°C for 30 seconds, 72°C for 20 seconds and final termination at 72°C for 5 min in a thermocycler (Applied Biosystems, USA). All PCR products were resolved and visualized by 1.5% to 2% w/v agarose gel electrophoresis. For DNA sequencing, the specific band was gel-excised and gel-extracted using QIAQuick Gel Extraction Kit (Qiagen, Germany). The eluted DNA concentrations were then measured by the NanoDrop spectrophotometer (Thermo Fisher Scientific, UK). The PCR products were subcloned into a TOPO TA cloning vector (PCRTM4-TOPO-Invitrogen TA cloning kit, Thermo Fisher Scientific, UK) and transformed into XL-10 Gold ultracompetent cells (Agilent Technologies, USA). Two clones bearing inserts of each sample were sent for Sanger Sequencing to Genewiz, UK.

#### Global Methylation

Global methylation was measured using a colourimetric, ELISA adapted method, MethylFlash Methylated DNA 5-mC Quantification Kit (Epigentek, US) as per manufacturer’s instructions. DNA (5 ng) was used for quantification and absorbance (405nm) was read using the Molecular Devices SPECTRA MAX PLUS Plate Reader Ver. 3.05 Spectrophotometer.

#### Whole-genome bisulphite sequencing

Whole-genome sequencing of uninfected and *T. gondii-infected* PC12 cells was performed at the University of Leeds Next Generation Sequencing facility at St James Hospital. DNA samples were collected by phenol-chloroform extraction as previously described. Five biological replicates were pooled for uninfected and control samples. Libraries were prepared using the TruSeq DNA Methylation Kit (Illumina, UK). Briefly, samples were bisulphite treated using the EZ DNA Methylation-Gold Kit (Zymo). Treated DNA was then amplified and 5’ adapters ligated. 3’ indexing adapters were then ligated to the ssDNA and 12 rounds of PCR performed. Sequencing was performed using the HiSeq 4000 Systems (Illumina). Differential methylation between infected and uninfected samples was calculated based on genome alignments by the Facility.

#### Statistical Analysis

All statistical analysis was performed using GraphPad and SPSS. For each data set, Levene’s test of equal variance was performed. Unless otherwise stated, a Student’s t test and where appropriate ANOVA with a post hoc test were performed for all equally distributed data.

## Acknowledgements

The authors wish to thank Mohammad Alsaad and Steve Clapcote who performed mouse infections, dissection and cryostat sectioning, and Greg Bristow who performed rat infections and tissue collection. We thank Dong Xia for the proteomic analysis of the EVs. Additionally, Martin Fuller for electron microscopy and Agne Antanaviciute performed Methylseq genome analysis. FACS was performed with the assistance of Sally Boxall at the Cell Sorting Facility of the University of Leeds.

## Ethics Declaration

The authors declare no competing interests.

## SUPPLEMENTAL INFORMATION

**Figure S1:**
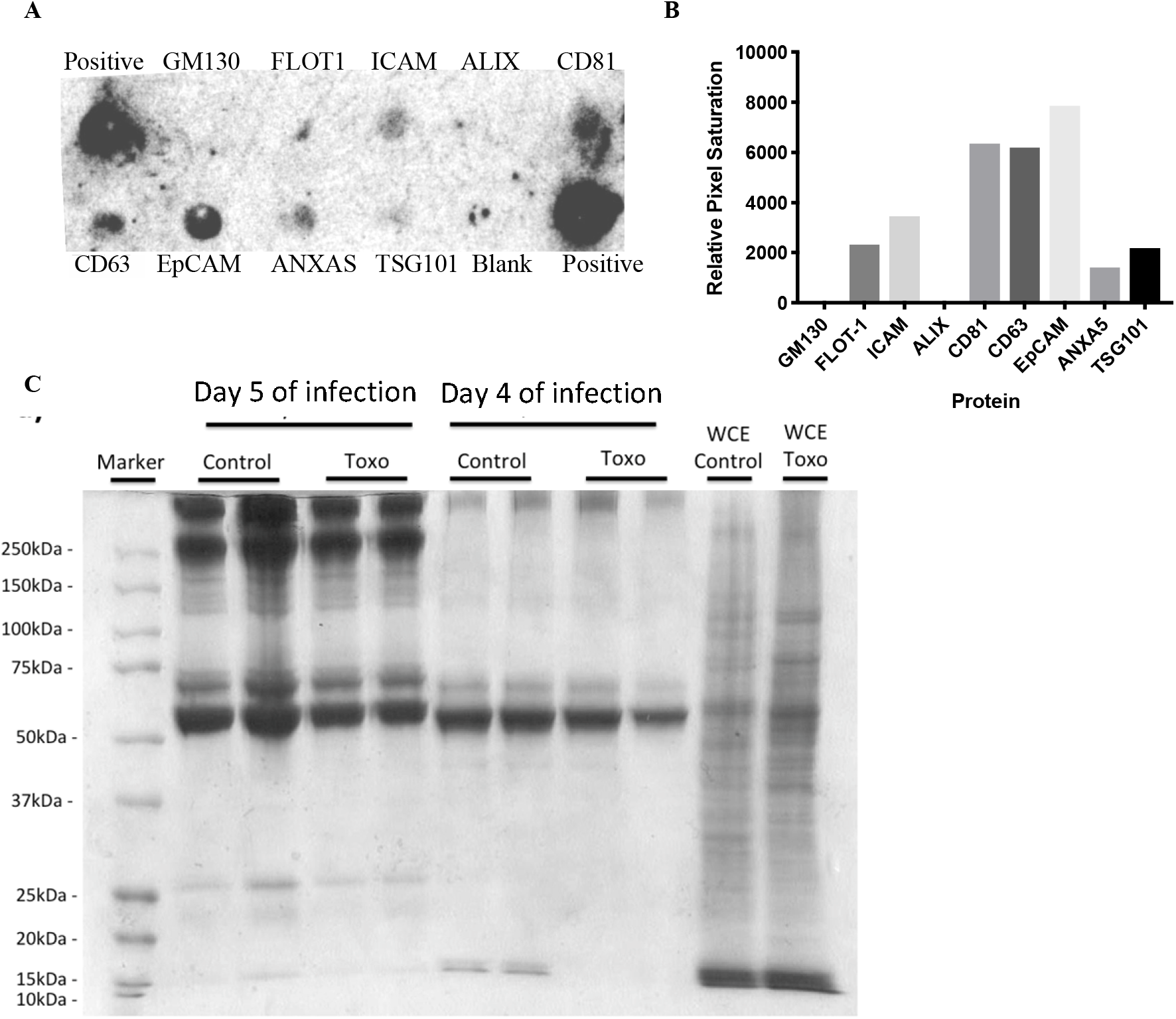
Extracellular vesicles isolated from infected cells exhibit classical exosomal markers. **A)** Image of a commercial dot blot probed with protein (300μg) from purified extracellular vesicles from *T. gondii* infected rat catecholaminergic cells. The dot blot contained eight common exosome markers and GM130 to indicate cellular contamination. (Exo-Check™ Exosome Antibody Array, System Biosciences, Palo Alto) with upper left and lower right HRP-conjugated antibody as positive controls. **B)** Quantitation of luminescence in the blot of the exosome markers using image J software. **C)** Protein analysis of EVs isolated from PC12 cells and infected cells. Purified protein from EVs was resolved by 12% SDS-PAGE and a coomassie blue stained image is shown. Protein markers and control whole cell extracts (WCE) are shown.

**Figure S2:**
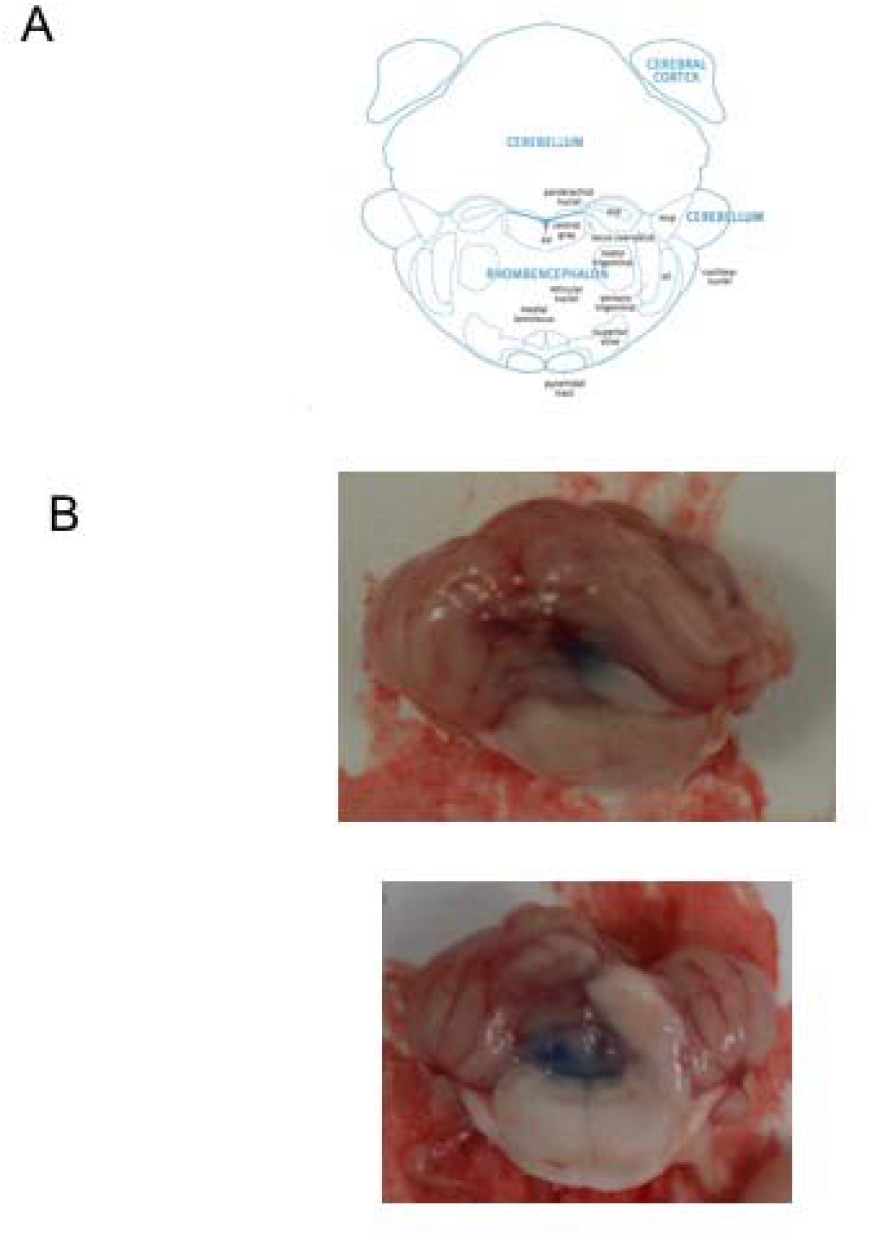
Delivery of EVs to the noradrenergic center, the locus coeruleus. **A)** Diagram of coronal section of the rat hindbrain taken from Paxinos and Watson, Academic Press **The Rat Brain in Stereotaxic Coordinates 6^th^ ed. B)** Examples of hindbrains dissected from rats shows localization of EV delivery by the bromophenol blue dye delivered through the cannula.

**Figure S3:**
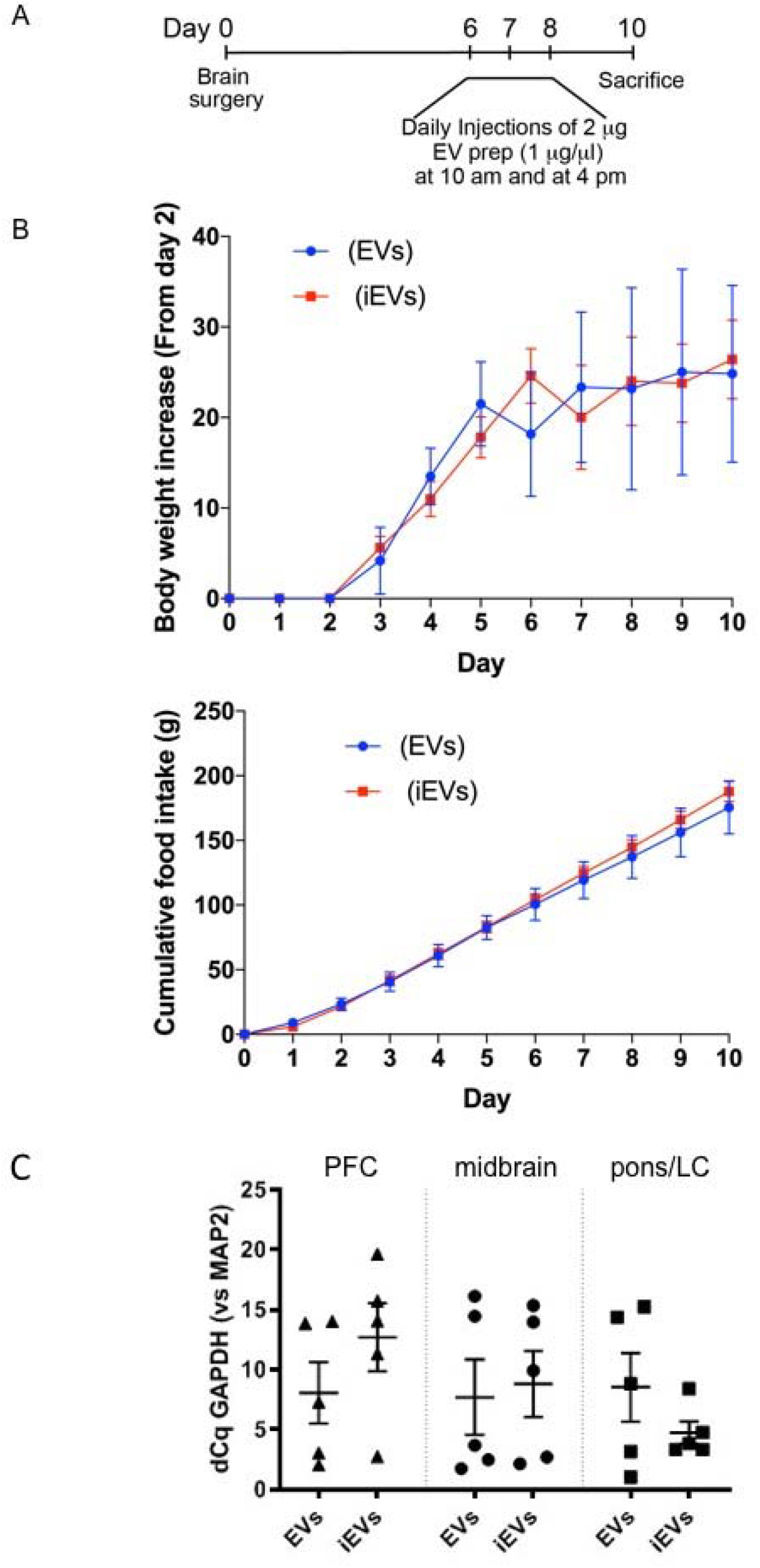
Intracerebral injection of extracellular vesicles (EVs) effect on health status and brain GAPDH expression. **A)** Timeline of the protocol for intracranial treatment of rats with purified EVs **B)** Body weight and appetite, measured as food consumption, in animals; during the course of the experiment. Following recovery from surgery these are normal. C) Expression of GAPDH in brain cells following EV treatment. Rats received *T. gondii-induced* neuronal extracellular vesicles (TINEV) labelled as iEV or EVs purified from uninfected cultures. Brains were sectioned into the prefrontal cortex (PFC), midbrain, and the hindbrain containing the locus coeruleus (pons/LC). Expression of GAPDH, as a representative gene for numbers of astrocytes, oligodendrocytes, microglia, and neurons shown relative to the neuronal marker MAP2. dCq is difference in Cq quantification cycle between GAPDH and MAP2, ±SEM shown, n=5, Student’s t test comparing TINEVs and control EVs for the regions: p = 0.86, 0.28, and 0.26, respectively.

**Figure S4:**
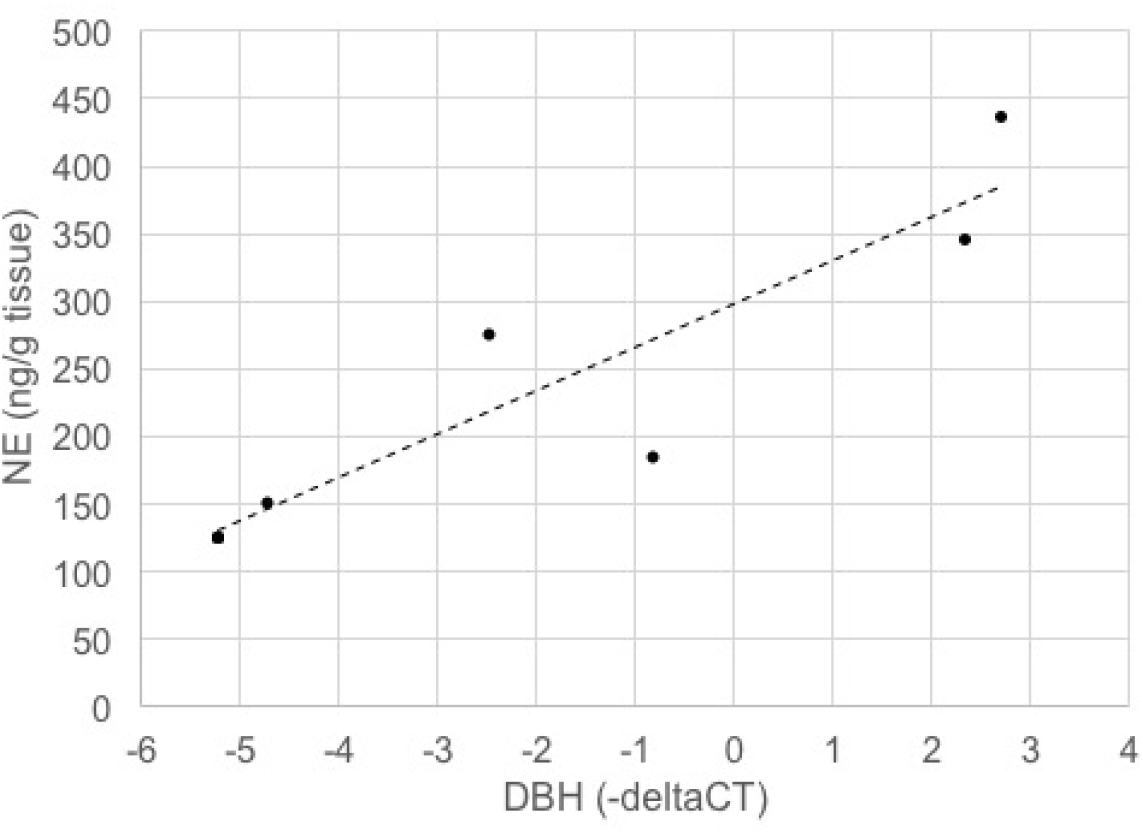
Dopamine β-hydroxylase during infection directly correlates with norepinephrine (NE) levels in the brain. Norepinephrine levels were measured by HPLC-ED and dopamine β-hydroxylase (DBH) was quantitated by RT-qPCR (relative to GAPDH) in brain tissue from rats. The Pearson’s correlation coefficient is 0.81 and the R value is 0.90 (p=0.014).

**Figure S5:**
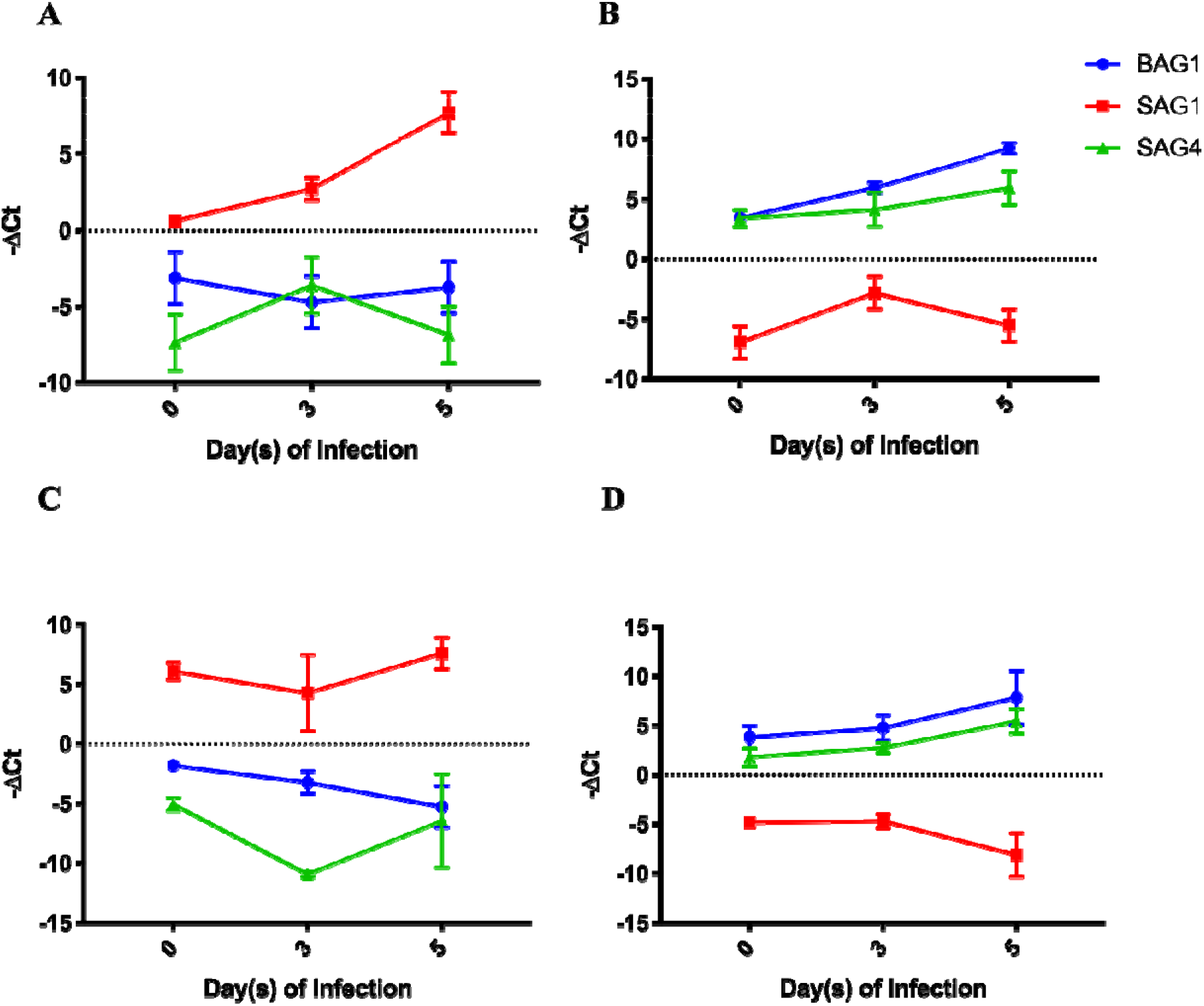
pH shocking of liberated *T. gondii* induced a bradyzoite-like phenotype. All plots show expression of the bradyzoite markers BAG1 (blue) and SAG4 (green), as well as the tachyzoite marker SAG1 (red) expressed in relation to the housekeeping gene GAPDH over a time course of 5 days of infection. **A)** Standard infection with tachyzoites with RNA collected over a time course of 5 days post-infection of rat catecholaminergic PC12 cells. ±SEM shown, n=6. B) Infection of Infection with pH-shocked *T. gondii*, as described in the Methods, of PC12 cells and RNA harvesting for RT-qPCR over a 5 days time course. ±SEM shown, n=6. **C)** Standard infection with tachyzoites as in A of human neuronal M17 cells and sample collection and processing. ±SEM shown, n=6. D) Human neuronal M17 cells were infected with pH shocked *T. gondii* and cultured for 5 days. RNA was collected on day(s) 0, 3 and 5; RT-qPCR was then performed ±SEM shown,, n=6.

**Figure S6:**
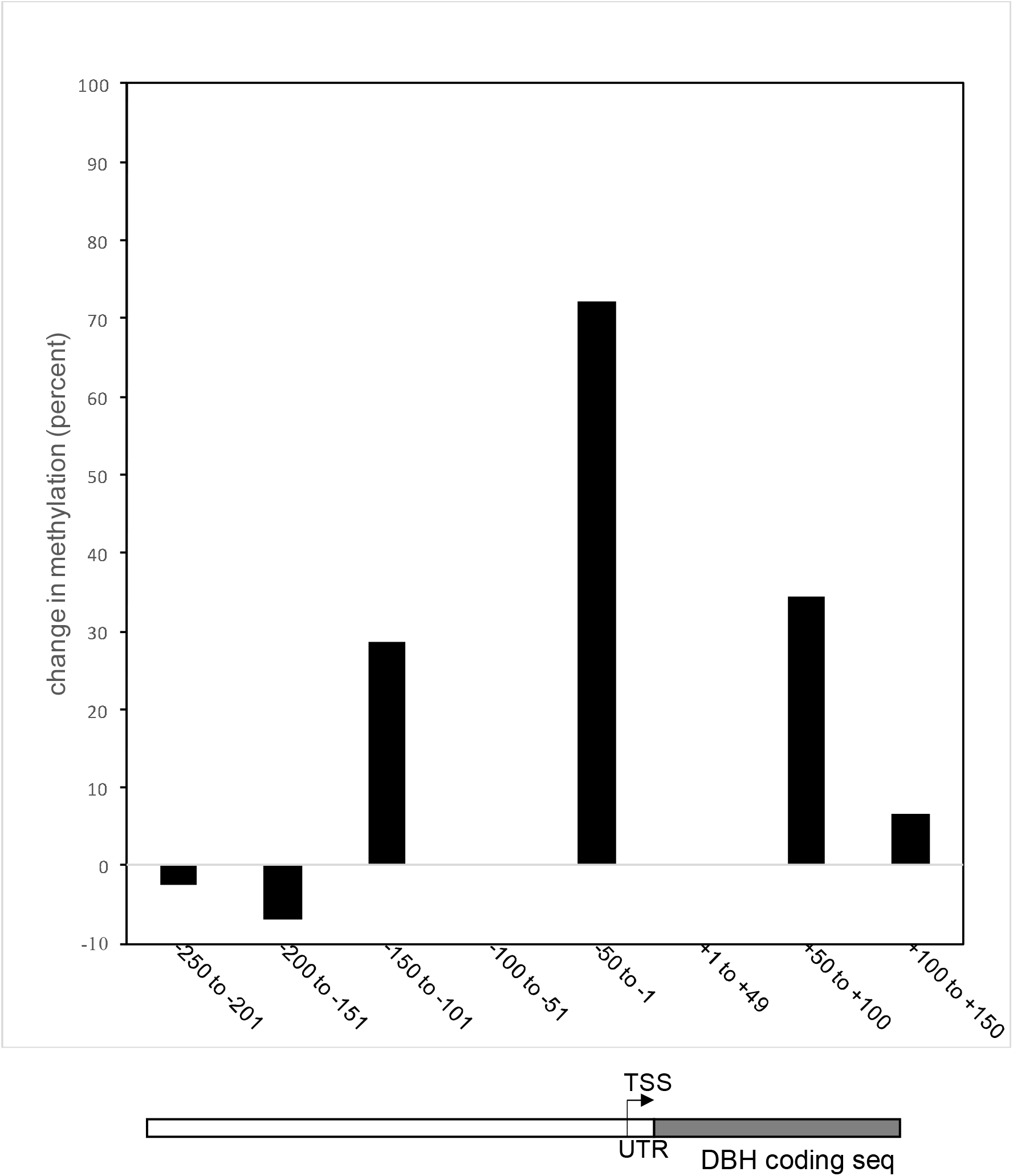
Methylation in the DBH upstream region based on bisulfite sequencing. Graph depicting the changes in CpG methylation across the *DBH* gene region from massively parallel genome sequencing for infected noradrenergic PC12 cells from three infected and three uninfected cultures at day five of *T. gondii* infection aligned with a diagram of the gene.

**Figure S7:**
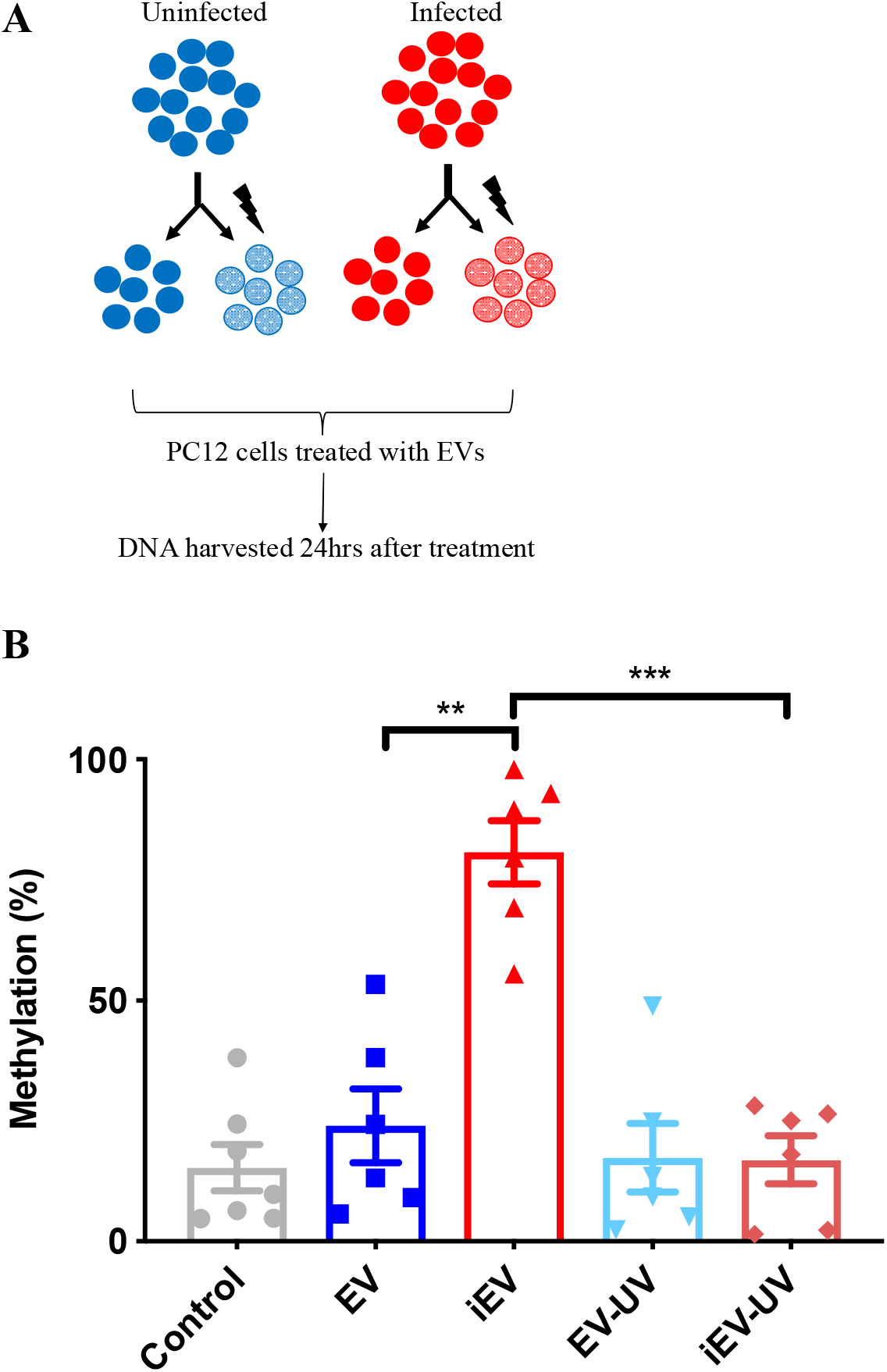
UV irradiation reverses the effect of TINEVs on 5’ promoter methylation of *DBH.* **A)** A schematic representation of experimental design. TINEVs and EVs derived from PC12 cells were mock or UV irradiated for 10 minutes at 4°C. These were used to treat cells for three days followed by DNA harvesting and methylation analysis. B) Methylation of the 5’ region of *DBH*, measured by MSRE qPCR in treated cells. Samples were from untreated cells untreated (grey), cells treated with EVs from uninfected cells (dark blue), TINEVs (labelled iEV; red), UV-irradiated EVs from uninfected cells (light blue), and UV-irradiated TINEVs (dark red); n=6 ±SEM shown. Two-way ANOVA performed, ** p<0.01, *** p<0.001. Tukey’s post hoc test for TINEVs compared to controls and UV treatments, p=0.001; and between controls and UV treatments, p= 0.90.

**Figure S8:**
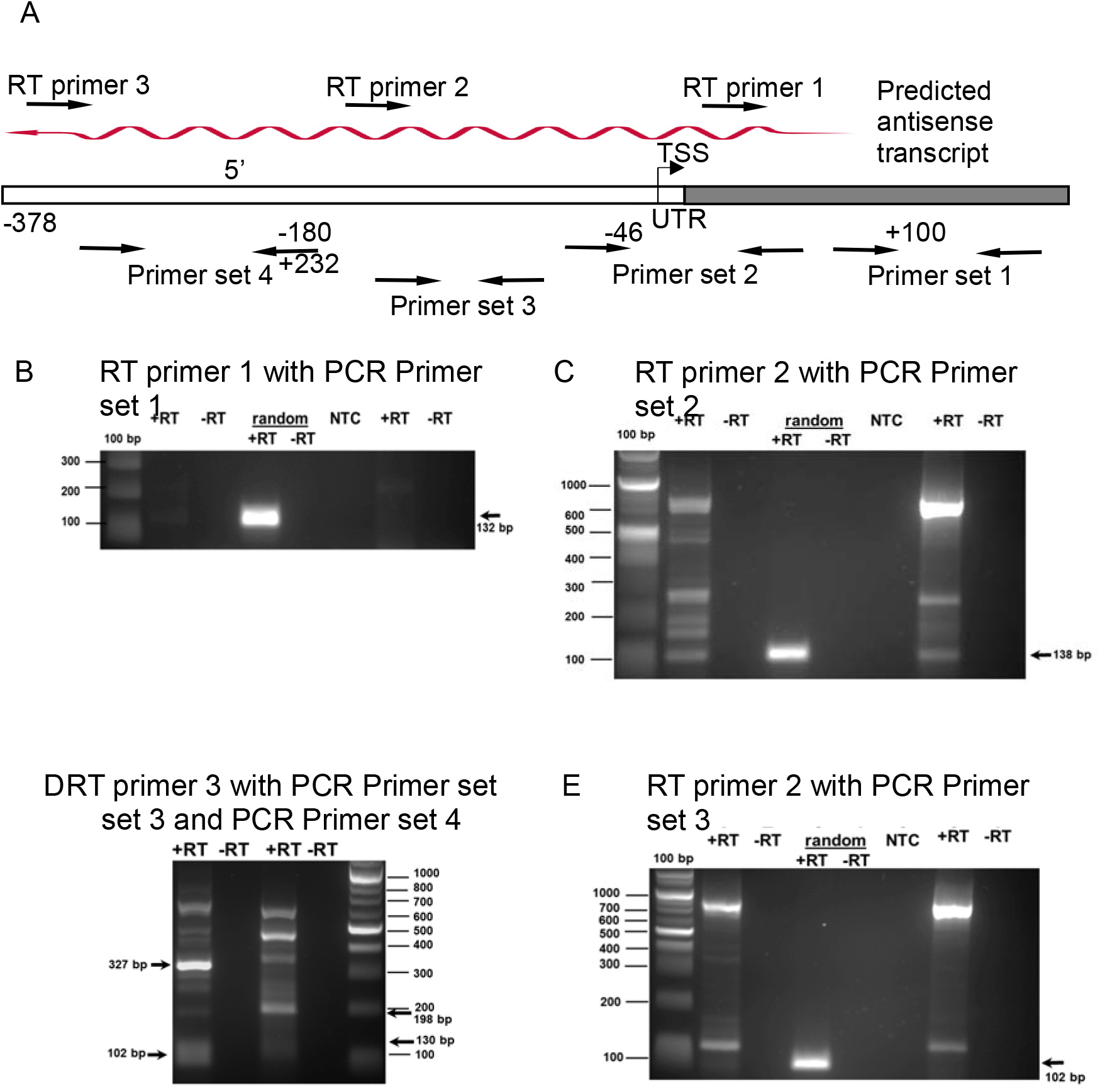
Genome scanning for DBH antisense lncRNA. **A)** Schematic representation of the stepwise walking upstream of the *DBH* gene to identify a putative antisense lncRNA. The binding sites for the directional RT primers for antisense transcript detection are depicted. Primer sets 1-4 were used to amplify cDNAs produced by the directional primers as detailed below. The transcription start site (TSS), untranslated region (UTR) and protein coding region (grey) are highlighted. **B-E)** *DBH* antisense RNA detected in *T. gondii-infected* PC12 cells. Agarose gel resolution of PCR products from directional reverse transcriptase (RT) primers. Control reactions lacking reverse transcriptase (-RT) were performed for each sample. Random hexamer primers that will detect both + and – strands in reverse transcription reactions were also tested. B) antisense *DBH* was not detected as a PCR product (132 bp) using directional RT primer 1 (+13) and PCR primer set 1 (+100 to +232) although the random hexamer primers yielded a band indicating *DBH* mRNA. C) antisense *DBH* from directional RT primer 2 (−180) was detected by PCR primer set 2 (−46 to +92) as a PCR product (138 bp). Additional bands are visible. **D)** Directional RT primer 3 (−378) detected antisense *DBH* with amplification by PCR primer set 3 (−153 to −51) with a PCR product (102 bp) and by PCR primer set 4 (−310 to −180) with 130 bp PCR product. Unexpected bands at 327 bp and 198 bp are due to amplification using the RT primer 3 and reverse primers for PCR primer set 3 and 4, respectively. E) First strand synthesis with RT primer 2 and PCR with primer set 3 yielded an unexpected band at 129 bp rather than the predicted 102 bp band. This was due to PCR amplification utilizing RT primer 2 and the reverse primer in set 3. PCR products of bands on gels were confirmed by DNA sequencing.

**Figure S9:**
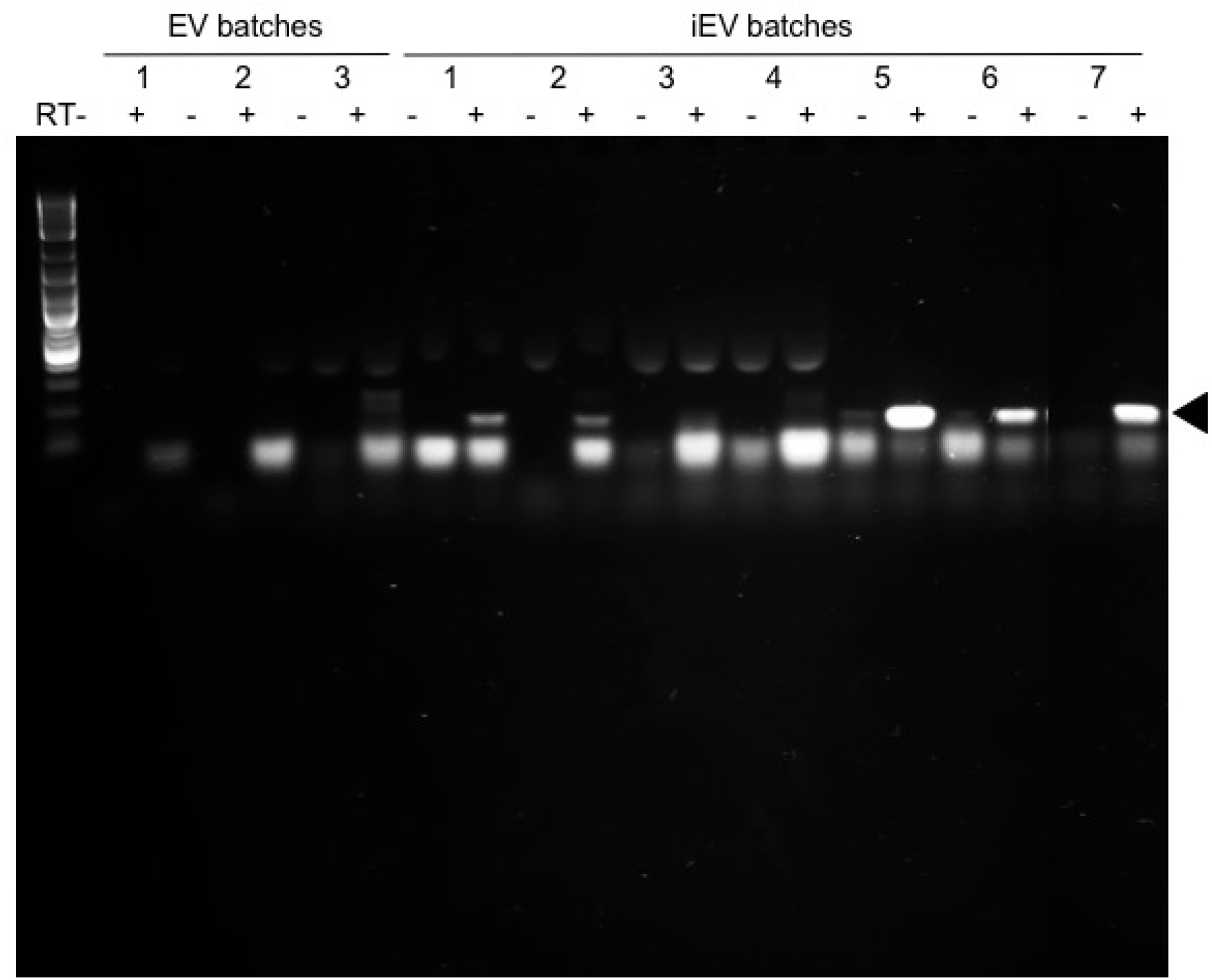
Agarose gel image of antisense *DBH* detection in TINEVs. The RT-PCR products resolved on a 1.5% agarose:TAE gel as in Fig 6D are shown with the full gel image. Antisense *DBH* from EVs from uninfected PC12 cultures (n=3) and TINEVs preparations (n=7) are shown with alternating lanes lacking reverse transcriptase. The PCR product (138 bp) is denoted by an arrowhead. The product was verified by cloning and sequencing. The first lane is a DNA ladder (NEB 1 kb plus). A lower (≈100 bp) background band appeared in some lanes.

### SUPPLEMENTAL DATA TABLE

**1:**
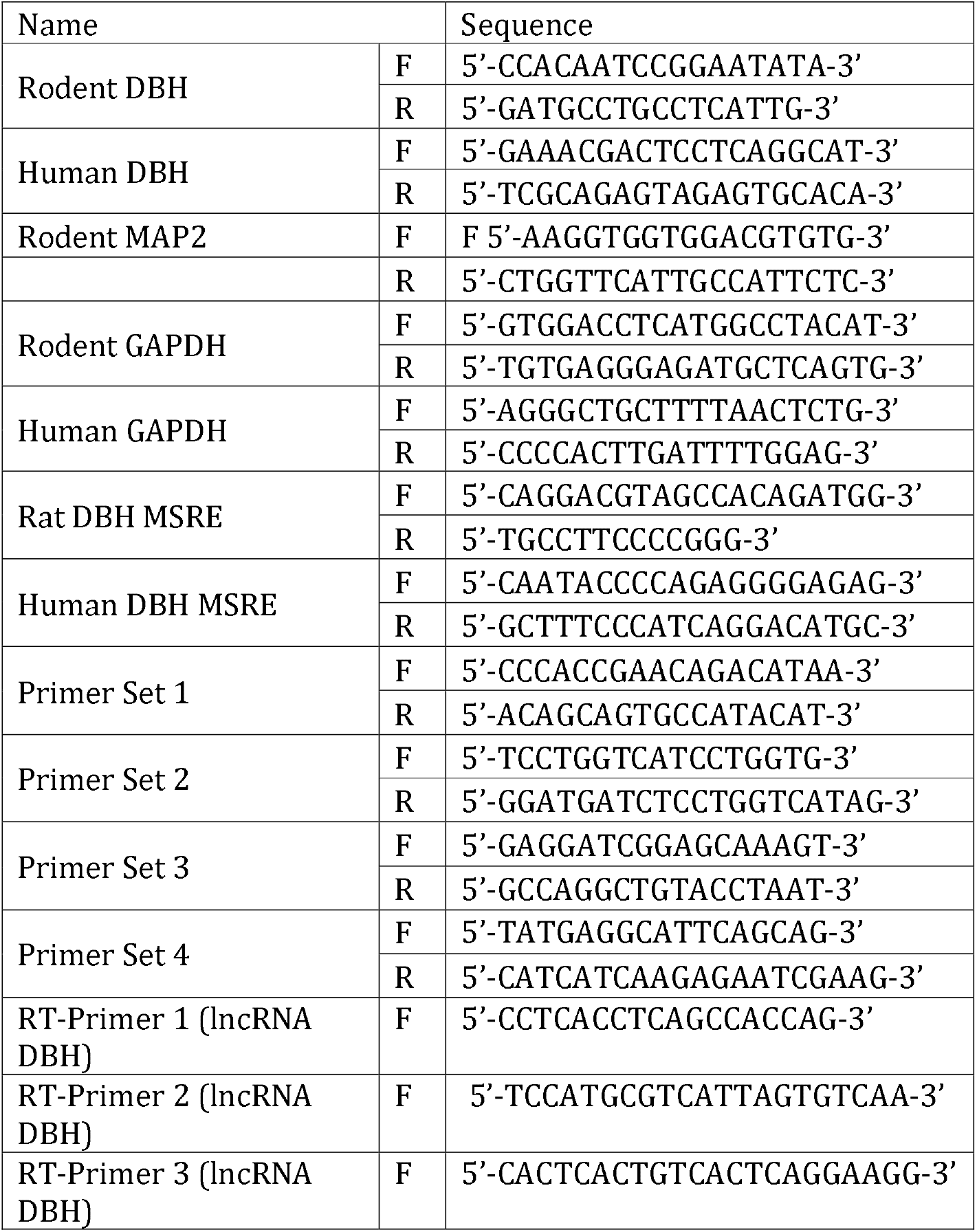
Primer sequences used throughout.

